# The short conserved region-2 of LARP4 interacts with ribosome-associated RACK1 and promotes translation

**DOI:** 10.1101/2024.11.01.621267

**Authors:** Amitabh Ranjan, Sandy Mattijssen, Nithin Charlly, Isabel Cruz Gallardo, Leah F. Pitman, Jennifer C. Coleman, Maria R. Conte, Richard J. Maraia

## Abstract

LARP4 interacts with poly(A)-binding protein (PABP) to protect mRNAs from deadenylation and decay, and recent data indicate it can direct the translation of functionally related mRNA subsets. LARP4 was known to bind RACK1, a ribosome-associated protein, although the specific regions involved, and relevance had been undetermined. Here, yeast two-hybrid domain mapping followed by other methods identified positions 615-625 in conserved region-2 (CR2) of LARP4 (and LARP4B) as directly binding RACK1 region 200-317. Consistent with these results, AlphaFold2-multimer predicted high confidence interaction of CR2 with RACK1 propellers 5-6. CR2 mutations strongly decreased LARP4 association with cellular RACK1 and ribosomes by multiple assays, whereas less effect was observed for PABP association, consistent with independent interactions. CR2 mutations decreased LARP4 ability to optimally stabilize a β-globin mRNA reporter containing an AU-rich element (ARE) more significantly than a β-globin and other reporters lacking this element. While polysome profiles indicate the β-glo-ARE mRNA is inefficiently translated, consistent with published data, we show that LARP4 increases its translation whereas the LARP4-CR2 mutant is impaired. Analysis of nanoLuc-ARE mRNA for production of luciferase activity confirmed LARP4 promotes translation efficiency while CR2 mutations are disabling. Thus, LARP4 CR2-mediated interaction with RACK1 can promote translational efficiency of some mRNAs.

## INTRODUCTION

Gene expression includes intricate control of mRNA stability and translation (1). Deadenylation of mRNA poly(A) tails (PATs) is linked to eukaryotic mRNA decay, occurs cotranslationally, and is modulated by the cytoplasmic poly(A)-binding protein (PABPC1/PABP) and associated proteins (2). In addition to binding poly(A), PABP functionally interacts with translation initiation factors (3–6). mRNA PAT length is coupled over a wide range to translational efficiency (TE) during embryogenesis, dependent on limiting PABP levels (7,8), while in other cells in which PABP is higher (8,9), the range in TE is substantially diminished (see 10).

Five families of La-related proteins (LARPs) share a La-motif (LaM) RNA binding domain (11,12), which in most members is followed by a family-specific RNA recognition motif (RRM) (for exceptions see 13,14). LARP4 and LARP1 have additional family-specific regions that have been shown to interact with PABP (15–19). LARP founding member, the authentic nuclear La protein (LARP3) uses conserved amino acids in its LaM aided by an edge of the RRM to recognize and sequester the U(2–3)U-3’OH motifs shared by RNA polymerase III transcripts (20,21), which protects them from 3’-exonuclease activities (20,21) (reviewed in 12).

The LARP1-PABP complex impedes global mRNA deadenylation by protecting short length PATs from 3’-exonuclease activity of the CCR4-NOT (CNOT) deadenylase complex (19). LARP4 also hinders mRNA deadenylation although at short PATs distinct from LARP1 and at long PATs (15,19). Results from cells deleted of *LARP4* and expressed at multiple levels revealed that LARP4 (aka LARP4A) slows deadenylation globally, targeting long and short PATs, the latter of which promotes ribosomal protein (rp) mRNA stabilization (15,22). The LARP1 family-specific DM15 domain binds the m**^7^**G-capC 5’terminal oligopyrimidine (TOP) motifs of rp-mRNAs to regulate their translation and stability (23–25). Crystal structures showed the conserved LaM of LARP1 recognizes A(2–3)A-3’OH, which PAT-seq confirmed confers cellular mRNA PAT protection-stabilization by LARP1 (13). The LaM and adjacent PABP-interaction motif, PAM2 of LARP1 confers mRNA PAT protection-stabilization as a small protein fragment lacking DM15, likely reflecting global activity (17).

As LARP4-PABP interactions are examined in this study, a brief overview follows. PAM2 is a consensus sequence found in ∼20 proteins involved in mRNA metabolism (18,26) that competitively bind the C-terminal MLLE domain of PABP (18,26,27). The PAM2w sequences unique to LARP4 and its paralog LARP4B/LARP5, present tryptophan (W) at consensus position-10 replacing phenylalanine in other PAM2s (18,28–30). The LARP4 PAM2w serves as a sequence-specific determinant of binding to either poly(A) or to MLLE (31). While less is known about the other PABP-binding motif (PBM) in LARP4 (and 4B), some data suggest it is more critical for PABP interaction and for mRNA PAT protection-stabilization than the PAM2w (22,28).

The family-specific PAM2 and PBM appeared in vertebrate LARP4B and LARP4/4A lineages along with conserved region-2 (CR2) in their less overall-conserved C-terminal half (32) (**Fig 1A**). Although LARP4 and LARP4B are distinguished by functional specifications (33), they share mRNA PAT-protection activity, while La, LARP6 and LARP7 do not (22). Recent data suggest they also share an apparent mRNA subset-specific, translation activity at the mitochondrial outer surface, likely distinct from LARP4 general activity (34). Indeed, LARP4 is recruited with PABP by A-kinase anchoring protein-1 (AKAP1) which direct mRNAs encoding mitochondrial proteins to mitochondria (35) (see 36). LARP4B and LARP4A each cofractionate and colocalize with TOM20 at mitochondrial outer surfaces suggesting this is a deep-rooted, shared function (37).

**Figure 1.**
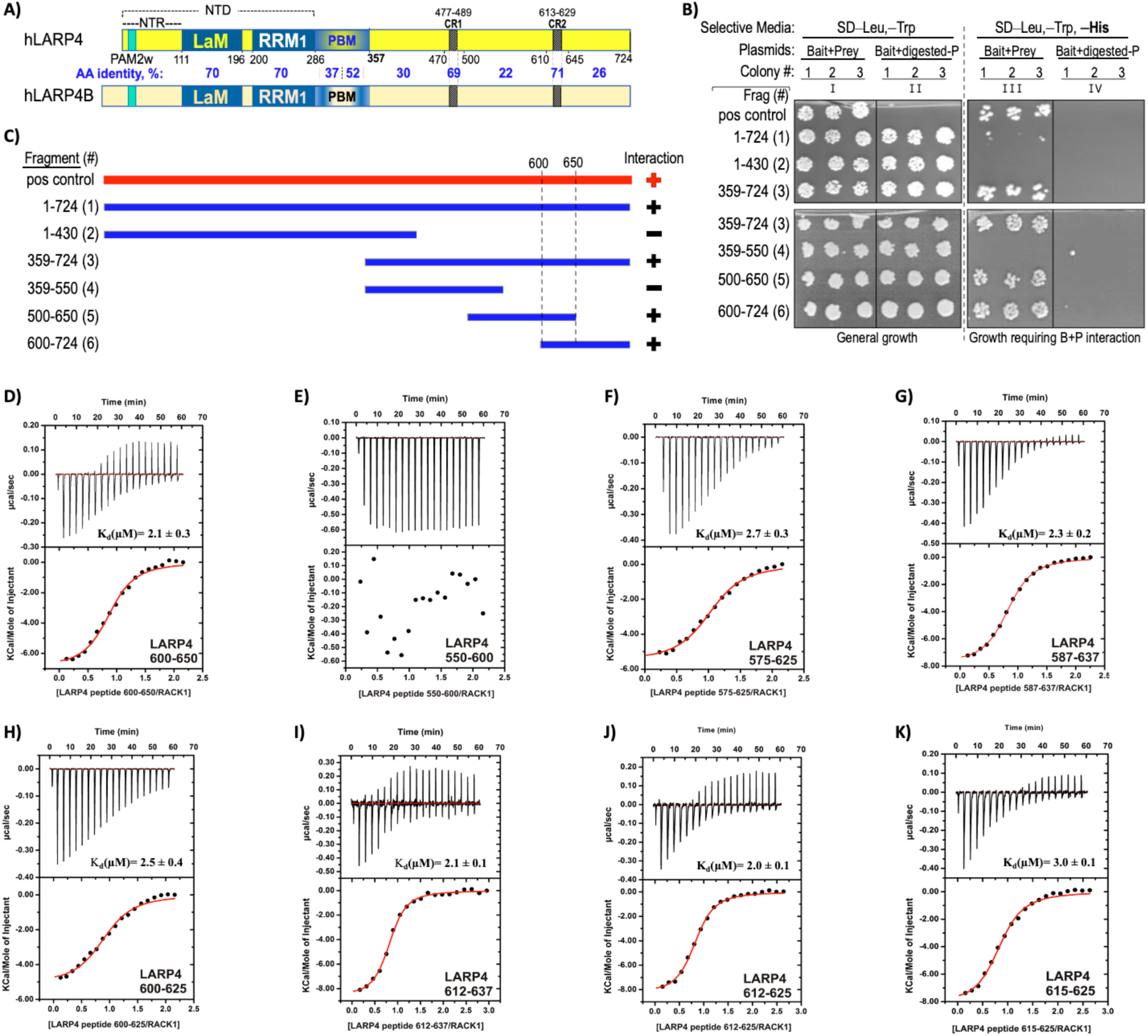
Mapping a minimal LARP4 region that interacts with the RACK1 region encoding propeller blades 5-7. **A)** Schematic representation of human LARP4(1–724) with amino acid positions demarcated below the boundaries of its known motifs and domains by black numbers. NTR: N-terminal region, PAM2w: PABP-interacting motif*-*2, w-variant (see text), LaM:La Motif, RRM1: RNA recognition motif-1, PBM: PABP-binding motif, CR1: conserved region 1 (residues 477-489), CR2: conserved region 2 (613–629). The bracket above indicates the NTD comprised of residues 1-286. Schematic of the human paralog LARP4B is shown below; amino acid (AA) sequence percent identity of domains and other regions between demarcated boundaries are in blue font. **B)** Yeast two hybrid (Y2H) domain mapping results obtained with the six LARP4 bait clones used as numbered fragments in panel (C) along with the full-length LARP4 positive control (“pos control”). The Frag #s in parentheses and their domain boundaries are indicated to the left of column I. Each column shows growth of three colonies (colony #) after liquid growth as indicated. The selective media in the agar plates is indicated on the top line. Prior to transformation the prey plasmid was either digested with a linearizing restriction enzyme (columns II and IV) or not (I and III). Growth on selective plates in column III reflects interaction with RACK1(200–317). **C)** The six LARP4 fragment clones used in Y2H assay shown as blue horizontal bars, and the full-length LARP4 positive control shown in red as bait, with RACK1 ORF200-317 as prey Positive and negative interactions are indicated by plus and minus respectively. **D-K)** Isothermal titration calorimetric analysis (ITC) of association between synthetic peptides corresponding to LARP4 fragments and recombinant full-length RACK1 protein. For each graph, the upper panel corresponds to the raw titration data showing the thermal effect of injecting solutions of the LARP4 peptide into a calorimetric cell containing RACK1 protein. The bottom panel shows the binding isotherms created by plotting the integrated raw data against the molar ratio of the peptides. The *K***_d_** and other thermodynamic parameters derived from this analysis (in triplicate) are reported in Table 1.

Studies of LARP4 mechanism use model mRNA reporters that contain or lack the TNFα ARE (13,17,22), a high-affinity binding site for TTP/ZFP36 (38) which can recruit CNOT deadenylase complexes by two of the several subunits (39–41). Classic stable mRNA reporters which lack an ARE and bear long PATs are substrates of the PAN2-PAN3 deadenylase, which is recruited to PABP by a PAM2 on PAN3 (27). In human cells, PAN2-PAN3 gradually shorten long PATs, unlinked to mRNA decay, to <120 nt, at which they become substrates of CNOT activity which is linked to their decay (42). By recruiting CNOT via an adapter protein, ARE-mRNAs bypass slow deadenylation and succumb to fast decay (2). Intriguingly by contrast, short PAT transcripts such as stable rp-mRNAs appear to sustain efficient translation as CNOT substrates (43) (see 2). Notably, LARP4 activity to slow deadenylation extends to stable transcripts, including rp-mRNAs, manifested with lengthened PATs and increased abundance (15,22).

RACK1 is a seven-propeller WD40 repeat protein that stably binds 40S subunits on ribosomes (44). RACK1 is multifunctional, participates in resolution of stalled, collided ribosomes and the associated stress response (45–48), and is a hub for the integration of factors that access translating mRNAs (49). Though RACK1 interacts with LARP4 and LARP4B, a binding region had been identified only for the latter, broadly within the >350 residues from the end of RRM1 to the C-terminus, and consistent with this location was shown to be noncompetitive with PABP binding (50). Thus, we sought to refine and examine this interaction further.

Yeast two hybrid (Y2H) domain mapping, biochemical binding assays, coimmunoprecipitation (co-IP), and cosedimentation identified LARP4 CR2 residues 615-625, including a central highly conserved SYAEVC sequence as a principal binding site for RACK1 region 200-317. Consistent with this, AlphaFold2-Multimer (AFM) predicted high confidence interaction between the LARP4 CR2 and RACK1 propellers 5-6. CR2 mutations disabled direct interaction with RACK1 in binding assays as well as by cosedimentation with ribosomes and polysomes. The CR2 mutations also impaired LARP4 for optimal stabilization of a reporter β-globin-ARE mRNA more significantly than impaired stabilization of β-globin mRNA and other reporters lacking the ARE. Translational deficiency was associated with presence of the ARE in β-globin mRNA, but not β-globin mRNA lacking the ARE, as monitored by polysome sedimentation profiles, was largely reversed by LARP4 while the LARP4-CR2 mutant was impaired. Analysis of nanoluciferase-ARE-mRNA by polysome sedimentation and luciferase activity production confirmed that LARP4 promotes its translation while the CR2 mutant is impaired.

## MATERIAL AND METHODS

### DNA constructs

LARP4 mutants (R1), containing five-point mutations R615E, K616E, Y619G, E621R, V622G and LARP4-ΔR1 (deletion of amino acids 612-625) cloned in the HindIII and BamH1 sites of pFlag-CMV2 were synthesized by Genewiz. The pcDNA3.1-β-Globin-TNFα-ARE and pcDNA3.1-β-Globin were subcloned from pTRERβ-TNF-α-ARE and pTRERβ-wt obtained from G. Wilson (51) as described (22); pcDNA3.1-TPGFP, obtained from J.R. Hogg (52), and previously characterized (13,15,17,22). LARP4B cDNA was subcloned from a LARP4B IMAGE clone (Source Bioscience) into a pCB6-GFP plasmid, using BamHI and XhoI restriction sites. To generate the LARP4B mutants, Site-Directed mutagenesis was performed. Mutagenic primers (Integrated DNA Technologies) were generated using NEBaseChanger™ online tool. Plasmid pCB6-GFP-LARP4B was mutagenized using the Q5 Site-Directed Mutagenesis Kit (NEB) according to manufacturers’ instructions. Plasmids were verified by sequencing (Sources Bioscience).

### Cell culture

PC3 cells (CRL-1435; ATCC) were cultured in RPMI 1640 (Gibco) with 10% FBS, 1% L-Glutamine (Sigma) and 1% penicillin/streptomycin (Sigma). Cells were kept in a humidified incubator at 37°C with 5% CO_2_ and passaged at 1:5 dilution every 3 days. HEK293 and HeLa Tet-Off cells (Clontech) were grown in DMEM with high glucose containing Glutamax (ThermoFisher, 10566016) and 10% FCS in a humidified incubator at 37°C with 5% CO_2_ and passaged at 1:10 dilution every 2 to 3 days.

### Yeast two-hybrid (Y2H) assays

Y2H assays were performed by Hybrigenics (Paris) to identify a LARP4 minimal region that interacts with RACK1 clone HLV_RP1_hgx1683v1_pB27_A-17 with ORF codons 200-317 obtained from a human liver cDNA screen (28). Human LARP4 and fragments thereof were used as bait.

### Isothermal Titration Calorimetry (ITC)

Human RACK1 protein was expressed and purified following the protocol described (53). HsLARP4 peptides 550-600, 575-625, 600-650, 587-637, 600-625, 612-667, 615-625 and 612-625 were ordered from GenScript (www.genscript.com). The lyophilized peptides were resuspended in MilliQ water to final concentrations ranging from 1 mM to 3 mM. The partially soluble peptide 600-625 was resuspended to 350 μM. Concentrations were measured by UV absorbance. ITC experiments were performed on an iTC200 microcalorimeter (Malvern Panalytical) at 298K in 20 mM HEPES, 300 mM NaCl, 10% glycerol and 1 mM DTT pH 7.5 (ITC buffer). The RACK1 protein and the peptides were diluted to the desired concentration in the ITC buffer. The peptide 600-625 (stock concentration 350 μM) was instead lyophilized and resuspended in the ITC buffer to avoid buffer mismatches. In each titration, volumes of 2µL of a 400 μM solution were injected into a solution of RACK1 (40 μM) with a computer-controlled microsyringe. For the experiments with peptide 600-625 and RACK1, the concentrations were 300 and 30 μM, respectively. Control titrations of the peptides into buffer alone were performed (not shown). The heat per injection normalized per mole of injectant versus molecular ratio was analyzed with the MicroCal-Origin 7.0 software package. ΔH (reaction enthalpy change in kcal/mol), K_b_ (binding constant equal to 1/K_d_), and n (molar ratio between the two proteins in the complex) were the fitting parameters. The reaction entropy was calculated using the equations Δ*G* = −*RTln K*_*b*_ and Δ*G* = Δ*H* − *T*Δ*S*. All ITC experiments were repeated at least three times.

### Transfection

Lipofectamine-2000 (Invitrogen) was used for transfections. HEK293 cells were seeded at 5.5 × 10^5^ per 6-well culture plate one day before transfection. The next day, 2.5 µg of pFlag-CMV2-LARP4-WT (15), mutant construct, or empty vector plasmid was transfected together with an aliquot of a pre-mixed batch of plasmids comprised of 400 ng of pcDNA3.1-β-glo-ARE, 50 ng of pcDNA3.1-β-Glo, and 100 ng pcDNA-TPGFP per 6-wells. Twenty-four hours after transfection, cells are split 1:3 and harvested 24 h later for protein and RNA. β-glo-ARE mRNA accumulates at several-fold lower levels than β-Glo mRNA. All experiments were planned and executed with caution regarding potential to saturate cellular ARE-mediated decay machinery with β-glo-ARE mRNA from transfected plasmid (54). Thus, amounts used are consistent with guidelines and notes on cell-specific variations, and our functional tests under conditions of actual experiments (below).

PC3 cells (3 × 10^5^ cells per 6-wells) were seeded 24 hr prior to transfection. Cells were transfected with 3µg of pCB6-GFP, pCB6-GFP-LARP4B and pCB6-GFP-LARP4B-R1^par^, 4.5µg of pCB6-GFP-LARP4B-R1, or 6µg of pCB6-GFP-LARP4B-ΔRIR.

### Northern blotting

Total RNA isolated using Tripure (Roche), was separated on 1.8% agarose-formaldehyde gels and transferred to GeneScreen-Plus membrane (PerkinElmer). After UV crosslinking and vacuum-baking for 2 hours at 80^0^C, membranes were prehybridized in hybridization solution (6 x SSC, 2x Denhardt’s, 0.5% SDS and 100 µg/ml yeast RNA) for one hour at the hybridization incubation temperature (Ti). **^32^**P-labeled oligo probes were added; hybridization was overnight at Ti.

### Immunoblotting

To isolate protein samples, cells were washed twice with PBS containing protease inhibitors (Roche) and cell lysis was performed directly in RIPA buffer (Thermo Scientific) containing protease inhibitors. Proteins were separated by SDS-PAGE and transferred to a nitrocellulose membrane. Primary antibodies (Ab) were anti-Flag-M2 (Sigma, F1804), anti-actin (Thermo Scientific, PA1-16890), anti-RACK1 (Santa Cruz Biotechnology, sc-10775), anti-LARP4B (Abcam, ab197085), anti-PABP1 (Cell Signaling, 4992S), anti-RPL9 (Abcam-ab182556) and anti-RPS6 (Abcam-ab225676). For all blots except anti-LARP4B, primary Abs were detected by secondary Abs from LI-COR Biosciences, which are conjugated to either IRDye 800CW or 680RD and the blots were scanned using the Odyssey CLx imaging system (LI-COR Biosciences). For anti-LARP4B, primary Ab was detected by HRP-conjugated secondary with detection by Clarity ECL substrates (Bio-Rad) and imaged using the ChemiDoc Touch system (Bio-Rad).

### Preparation of total cell extract for immunoprecipitation

HEK293 cells were transfected as above. 24 hours later; cells from duplicate wells were combined in a 10 cm^2^ culture dish. 24 hours later the cells were washed with 20 ml warm PBS, scraped in 10 ml ice-cold PBS, and transferred to a conical tube on ice. 5 ml ice-cold PBS was used to rinse the culture dish and added to the tube. The cells were spun in a pre-chilled centrifuge at 4^0^C for 3 mins at 1400 rpm. The supernatant was discarded, 200 μl lysis buffer added (50 mM Tris-HCl pH 7.5, 75 mM NaCl, 0.25% NP-40, 2x protease inhibitors (Roche), and made 2 mM PMSF). This was incubated for 5 mins on ice with occasional flicking of the tubes. After cell lysis was confirmed by inspection under the microscope, 600 μl ice-cold wash buffer (50 mM Tris-HCl pH 7.5, 75 mM NaCl and 0.05% NP-40) was added. Lysates were then spun for 15 mins at 13,000 rpm at 4°C. The supernatants were transferred to a new tube and their total protein concentration was determined by BCA (Pierce). A fraction of the clarified lysate from each sample was put at −80^0^C to be used as input and a fraction was used for IP. For LARP4B, PC3 cells were transfected with plasmids GFP-LARP4B, GFP-LARP4B-R1, GFP-LARP4B-R1**^par^**, GFP-LARP4B-DR1R, and GFP. Cells were pelleted 24 hours later, lysed, and the lysates were used for IP.

### Immunoprecipitation

For LARP4, 50 μl of anti-Flag M2 magnetic bead slurry (Sigma) was washed 3 times with 250 μl of wash buffer. HEK293 cell extract corresponding to 500 ug total protein was added to the beads in 340 μl total volume (adjusted by wash buffer 50mM Tris, pH-8.0; 75mM NaCl and 0.05% NP-40). Samples were incubated at 4°C for 2 hours, followed by 5 washes with 250 μl of wash buffer. The supernatant was removed, and beads suspended in SDS-PAGE sample buffer (without β-Mercaptoethanol, BME) followed by heating the beads at 95^0^C for 5 minutes and collecting the supernatant to which 5% fresh BME was added just before loading. These samples were heated to 95^0^C for 2 minutes and separated on SDS-PAGE gel followed by Immunoblotting. For immunoprecipitation of RACK1, 20 μl beads of Protein-A Sepharose were incubated with 10 μl of anti-RACK1 Ab, washed, and added to 500 μg HEK293 total cell protein extract. IP was as above. For LARP4B, PC3 cells were transfected with GFP vectors as described above. Cells were pelleted 24 hours after transfection. Cell pellets were lysed and the supernatant containing equal protein was diluted in 500 µl ice-cold wash buffer. Supernatants were added to pre-equilibrated GFP-TRAP Agarose beads (Chromotek) diluted in wash buffer and rotated end-over-end for 1 hour at 4°C. The beads were sedimented at 2,500 g for 5 min, the supernatant discarded and beads with IPed material washed three times in 500 µl wash buffer, before resuspension in 80 µl SDS-PAGE sample buffer (Laemmli, BioRad) containing 5% BME. Samples were heated at 95°C to disassociate complexes from beads. Beads were sedimented by centrifugation at 2,500 g for 5 min at 4°C and supernatants were analyzed as the bound fraction.

### Polysome profile analysis

Polysome fractionations were by standard methods (28,55) using a Brandel programmable density gradient fractionation system with UV detector (Foxy Jr.; Teledyne Isco, Lincoln, NE). Sucrose density gradients were made with a Gradient Master (Biocomp). HEK293 cells were seeded in 10 cm culture plates to achieve 80–85% confluence 16 h later. The cells were transfected with 8 µg or 16 µg of β-glo-ARE plasmid along with 800 ng of pcDNA3.1-β-Glo, 800 ng GFP plasmid and 20 ug pCMV2 plasmid containing either LARP4-WT or LARP4-R1. After transfection, the cells from each plate were split into two 15 cm culture plates. One day later, fresh sucrose solutions (47% and 7%, wt/vol) in 10 mM HEPES, pH 7.3, 150 mM KCl, 20 mM MgCl2, 1 mM DTT were prepared, filter sterilized and used to make the gradients. After a 5-minute incubation at 37°C after cycloheximide (Chx) (100 ug/ml) was added to the media, the plates were put on ice and washed twice with ice-cold PBS plus 100 ug/ml Chx. Five ml of ice-cold PBS with 100 mg/ml Chx was added per plate, the cells scraped and added to an ice-cold tube. The cell suspension was centrifuged at 1,200 rpm for 3 min at 4°C and the pellet taken up in 300 µl lysis buffer (10 mM HEPES, pH 7.3, 150 mM KCl, 20 mM MgCl2, 1 mM DTT, 2% NP-40, 2x EDTA-free protease inhibitors (Roche), 100 ug/ml Chx, and 40 U/ml RNaseOUT (Invitrogen)) and kept on ice for 2 min with occasional flicking. The lysate was cleared by centrifugation at 13,000 rpm for 5 min at 4°C. 400 µl was removed from the gradient tubes, and 10 OD260 absorbance units of lysate was carefully layered on top. The gradients were spun in an ultracentrifuge (Beckman SW41 rotor) at 33,000 rpm (134,000 g) for 2 hr and 50 min at 4°C. One ml fractions were collected; protein was precipitated from 250 µl by trichloroacetic acid (56); RNA was purified from 750 µl described below. Briefly, a concentrated lysate from the pool of transfected cells is placed atop and centrifuged through a 13-ml sucrose gradient during which free proteins and small complexes remain near the top, and 40S, 60S, 80S monosomes, disomes and polysomes sediment progressively deeper.

### S100 sedimentation through a sucrose cushion (57)

HEK293 cells were transfected with 2.5 µg of pCMV2 EV, LARP4-WT or LARP4-R1 and cultured as above. Forty-eight hours after transfection, cycloheximide was added to the media at 100 µg/ml, incubated at 37^0^C for 5 minutes and lysates prepared as described in the polysome section. OD260 absorbance equal to 60 µg RNA in 200 µl polysome lysis buffer was loaded on 800 µl of 1M sucrose cushion in 1 ml thick wall polycarbonate tubes (part numbers 343778; Beckman Coulter). After centrifugation at 90,000 RPM (352,000 x g) for 1h at 4^0^C in a TLA-100 fixed-angle rotor in Optima TLX Beckman ultracentrifuge, supernatants were collected. Pellets were dissolved in resuspension buffer (20 mM Tris-HCl, pH 7.6, 50 mM KCl and 5 mM MgCl_2_ with protease inhibitors) and precipitated with TCA. Input, supernatant and pellet fractions were immunoblotted. Quantifications were by the LiCor system.

### Quantification of mRNAs in polysome fractions

RNA was precipitated from 750 μl of polysome profile fractions by addition of 1.5 volumes (1125 µl) of RNA precipitation mix (95% ethanol, 5% sodium acetate pH 5.2), and incubated overnight at −20°C. Samples were centrifuged at 15,000 rpm for 30 min at 4°C and pellets washed with 1 mL cold 80% ethanol, air-dried, and resuspended in 100 µL of DEPC-treated H_2_O. RNAs were column purified using RNA Clean and Concentrator kit (Zymo R1018). The purified RNAs were separated on a 1.8% agarose-formaldehyde gel, transferred to a GeneScreen-Plus membrane (PerkinElmer), and stringently hybridized with 5’ **^32^**P labeled gene-specific oligo-DNA probes. For each specific mRNA all blots were hybridized over night with the same ^32^P probe, followed by stringent washing together, exposure together to the same PhosphorImager screen, and scanned using Amersham^TM^ Typhoon and quantified using ImageQuant. For each mRNA, the percent cpm in each fraction of total cpm in all fractions was plotted (58).

### β-glo-ARE mRNA half-life determinations

HeLa Tet-off cells were co-transfected with 100 ng pTRERb (β-glo-ARE under a Tet-responsive minimal CMV promoter) along with 2.5 µg of LARP-WT, -R1, -ΔPBM, or empty vector (EV) constructs. The next day, cells were equally divided into multiple wells. 48 hours post transfection, the media was replaced with fresh media containing 2 mg/ml doxycycline (Sigma) to block transcription from the Tet-responsive minimal CMV promoter and the cells were harvested at indicated times thereafter. Total RNA was analyzed by Northern blotting.

### Sequence analysis

Multiple sequence alignment (MSA) was derived from a MSA of the full-length proteins using MacVector. FASTA files used to create sequence LOGOs for CR2 of LARP4 and 4B were from the sequences in Deragon (32). FASTA files used for LOGOs of extended CR2 and CR1 were as follows; BLASTP https://blast.ncbi.nlm.nih.gov/Blast.cgi was used to query the NCBI nr-cluster database using 20 and 26 amino acid sequences respectively, with max target sequence parameter set to 500. For extended CR2, this is positions 613-632 of LARP4 which returned a wide variety of sequences annotated as LARPs 4, 4A and 4B. For extended CR1, two searches were done, one using LARP4 sequence positions 471-496 and the other using the corresponding region of LARP4B. The output results were downloaded from the MSA viewer, converted to FASTA files and uploaded to WebLogo-3 server at https://weblogo.threeplusone.com/create.cgi (59,60).

### AlphaFold-Multimer

was run using ColabFold (61) on UCSF ChimeraX (62) version 1.5. Protein sequences used as input sequences were: LARP4, NP_443111.4; LARP4B, NP_055970.1; RACK1, 317 aa, NP_006089.1; RPS3, residues 111-228 of genbank KJ892051.1; RPS17 residues 1-132 of NP_001012.1; uS9/RPS16, NP_001011.1. The exact input sequence regions of LARP4 and LARP4B used in different predictions varied and are therefore reported in the corresponding figure legends. Default ColabFold parameters were used to assess AFM prediction convergence, which includes 5 models per prediction, referred to as 5×1 from which a best model is selected (63). For random seed predictions, the “num_seeds=” was 5 which produces (5×5) 25 models or 25 (25×5) which produces 125 models, as stated in the text and/or figure legends.

### Pull-down assays with purified recombinant proteins

#### Plasmid construction

To generate pLIB-LARP4, a sequence encoding human LARP4 residues 1-724 was PCR amplified from a cDNA parent plasmid and inserted into pLIB (64) at the BamHI/HindIII restriction sites; primer-encoded C-terminal StrepII tag (65) was included in-frame with LARP4. pLIB-LARP4^Y619E^ and pLIB-LARP4^C623E^ were generated by site-directed mutagenesis of pLIB-LARP4 using overlapping primers LARP4(Y619E) _F/LARP4(Y619E) _R and LARP4(C623E)_F/LARP4(C623E) _R, respectively. Similarly, pLIB-LARP4^Δ478-488^ and pLIB-LARP4^Δ616-626^ were generated using overlapping primers LARP4(del478-488) _F/LARP4(del478-488) _R and LARP4(del616-626)_F/LARP4(del616-626) _R, respectively, and verified by sequencing.

To produce pnEK-NvH-RACK1, a sequence encoding human RACK1(1–317) was PCR amplified with primers HsRACK1_F/HsRACK1_R from a synthetic sequence codon-optimized for *E. coli* expression (Azenta). The amplified RACK1 sequence was inserted into pnEK-NvH (66), linearized with NdeI. The human PUM1-RD3 construct, residues 589-827, tagged with maltose-binding protein (MBP), was described (67,68).

#### Production and purification of human LARP4

Initial low-titer recombinant baculoviruses containing full-length human LARP4 under the control of a polyhedrin promoter were generated using transfection of adherent *Spodoptera frugiperda* Sf21 insect cells (Thermo Fischer Scientific) and then amplified in suspension culture at 27℃/120 rpm grown in SF900II serum-free media (Thermo Fisher Scientific). The higher-titer virus was used to infect Sf21 cells in suspension at a density of 2-million cells/ml, and cells were harvested 48 h after they stopped dividing. Cells were resuspended in lysis buffer (50 mM HEPES pH 8.0, 500 mM NaCl) and lysed using an Ultrasonics Sonifier SFX550 (Branson). Lysate was cleared by centrifugation at 40,000 *g* for 45 min at 4℃ and used immediately for immobilization on resin for pulldown experiments or stored at −20℃. Mutants and deletion variants were produced identically to WT protein.

#### Production and purification of RACK1

The human RACK1 was produced in *Escherichia coli* BL21 (DE3) Star cells (Thermo Fisher Scientific). The bacteria were grown in Luria-Bertani Miller medium (Research Products International) at 37℃ with shaking at 120 rpm until the optical density at 600 nm reached 0.3, at which point the temperature was reduced to 30℃. Expression was induced by adding 1 mM IPTG (GoldBio), and cells were harvested by centrifugation after four hours. The cells were resuspended in lysis buffer (50 mM HEPES pH 7.5, 1.0 M NaCl, 5% (v/v) glycerol, 25 mM imidazole) and lysed using an Ultrasonics Sonifier SFX550 (Branson). The lysate was then cleared by centrifugation at 40,000 *g* for 45 minutes at 4℃ The cleared lysate was then loaded on a 5 ml HisTrap FF column (Cytiva) at a flow rate of 2-5 ml/min using an Akta Pure chromatography system (Cytiva). Following the washing of the column, the bound protein was eluted over a linear gradient with elution buffer (50 mM HEPES pH 7.5, 200 mM NaCl, 5% (v/v) glycerol, 500 mM imidazole) over 10-20 column volumes at a flow rate of 2-5 ml/min. The protein was then diluted to 75 mM NaCl and loaded onto a 5 ml HiTrap Heparin HP (Cytiva) column at a flow rate of 2-5 ml/min. Following the washing of the column, the bound protein was eluted over a linear gradient of 75-1000 mM NaCl at a flow rate of 2-5 ml/min. Eluted fractions were analyzed for purity by SDS-PAGE followed by Coomassie staining and concentrated using Amicon centrifugal concentrators (Sigma-Aldrich). Concentrated protein was flash-frozen in liquid nitrogen and stored in 20-50 μl aliquots at −80℃.

#### Production and purification of MBP and MBP-tagged PUM1-RD3

These constructs were produced and purified as described (67,68). Briefly, bacteria were grown in auto-induction ZY medium at 37℃ for 16-18 hours with shaking at 120 rpm. The cells were resuspended in lysis buffer (50mM HEPES pH 8.0, 500mM NaCl, 5% (v/v) glycerol). The cleared lysate was prepared as above and used for pulldown experiments.

#### StrepII/StrepTactin pulldown assay

The procedure was described (69). Briefly, cleared lysates were incubated with 30 μl of StrepTactin Sepharose High-Performance resin (Cytiva) for one hour at 4℃. The resin was then washed 4-5 times with binding buffer (50 mM HEPES pH 8.0, 500 mM NaCl, 0.03% (v/v) Tween-20). 500 pmol of purified RACK1 protein was then added. After one hour of incubation, the beads were washed three times with binding buffer, and proteins were eluted with 50 mM biotin in the binding buffer. The eluted proteins were analyzed by SDS-PAGE followed by Coomassie staining. Each pulldown experiment was performed with at least two biological replicates.

### Generation of stable Flp-In-HEK293 NanoLuciferase expression cell lines

were established using the Flp-In™ System (Invitrogen). Different nLuc reporter constructs were cloned into the pCDNA5/FRT/TO plasmid, which contains the Hygromycin gene. These were then transfected into Flp-In HEK293 cells along with a Flp recombinase expression plasmid, pOG44. Following transfection and upon reaching ∼25% confluency, cells were split into 10 cm plates and subsequently subjected to selection by the addition of Hygromycin to the culture media at 100 µg/ml in Tetracycline (Tet) free media, which was refreshed every 2-3 days. As single colonies became visible after 1-2 weeks they were picked and propagated in 6-well plates in Hygromycin and Tet-free media. After 2-3 days of growth, cells from each well were split into two new wells. Following additional 24-hours, one well from each colony was induced with doxycycline at 1 µg/ml for 24 hours to activate the reporter gene expression. To the other well, an equal volume of sterile water was added to maintain the reporter gene in the uninduced state. After the 24 hr induction period, cells were harvested for protein and RNA extraction.

### Luciferase assays

were performed using a Thermo Ascent Luminometer which was replaced by a Promega GloMax® Navigator. Assays were prepared using Glo-Lysis buffer (Promega), supplemented with protease inhibitor. Protein concentrations of extracts were quantified using the BCA calorimetric assay following a standard curve generated using purified serum albumin. To achieve concentrations of cell extracts with RLU activity levels measurable within ∼100-fold of each other, the extracts were diluted to different extents, ranging from 1/10 to 1/40, in Glo lysis buffer containing 2 mg/ml BSA. Subsequently, 10 µL of each diluted sample was aliquoted into a white 96-well plate. 10 µL of Nano-Glo Luciferase Assay Reagent (1:50 mixture of Nano-Glo Luciferase assay substrate:Nano-Glo Luciferase assay buffer (Promega) was added to each well containing the diluted samples. The samples were thoroughly mixed in the 96-well plate by the shaking function of the luminometer, and luminescence was measured and recorded as relative light units (RLU) and where noted, normalized for each sample by the amount of protein, and expressed as RLU/µg of extract protein.

### LARP4 knock-out (KO) cells

were derived from the HEK293 Flp-In™ cell line (Invitrogen). LARP4 KO cells were confirmed to lack LARP4 expression via western blot. The cells were seeded at 7.0 × 10^5 per well in a 6-well plate one day before transfection. The next day, 2.5 µg of pFLAG-CMV2-LARP4 construct plasmids or empty plasmid (EV) was transfected with a pre-mixed batch of 1000 ng pcDNA3.1-β-Glo-TNFα−ARE and 100 ng pcDNA-TP-GFP plasmids per well. Lipofectamine 2000 was used at 2.0 µl per µg of DNA. Twenty-four hours post-transfection, the cells were split from one well into three (two for RNA extraction and one for protein extraction) and collected after another 24 hours, making it a total of 48 hours post-transfection.

## RESULTS

### Domain mapping of the LARP4 region necessary for binding RACK1 WD40 repeats 5-7

A Y2H screen using full-length (FL) LARP4 isolated four groups of clones encoding human RACK1 with different N-termini that shared open reading frame (ORF) codons 200-317 (28). Here, a RACK1 ORF200-317 bait clone, encoding propeller blades 5-7 (49) was used with a LARP4 domain mapping panel of six constructs to localize the interaction region. In addition to these, the original FL-LARP4 clone was a positive control (Fig 1B); further plasmid controls in appropriate selective media validated the Y2H assay (Supp FigS1). The FL-LARP4 construct and fragment 359-724 interacted, but 1-430 did not as in Fig 1B, column III consistent with C-terminal localized interaction LARP4B (50). As summarized in Fig 1C, further dissection revealed fragment #s 1, 3, 5 and 6 were positive for interaction while #s 2 and 4 were negative, revealing LARP4 positions 600-650 as the candidate Y2H domain mapping local region of interaction with RACK1(200–317).

Isothermal titration calorimetry (ITC) was used to examine the binding of full-length RACK1 with synthetic peptides representing LARP4. A peptide spanning amino acids 600-650 bound to RACK1 while the 550-600 peptide did not (Figs 1D-E). Several peptides that collectively spanned 575-637 (Fig 1F-I) showed that 575-625 and 587-637 bound to RACK1 with comparable affinity and a stoichiometry of 1, as did progressively shorter peptides 600-625, 612-637, 612-625 and 615-625 (Figs 1H-K). Thus, the minimal RACK1 interaction region tested was 615-625 (thermodynamic parameters are in Table 1).

**Table 1.** ITC thermodynamic parameters. (n is stoichiometry, K**_d_** is dissociation constant). Values are shown as average of triplicate measurements and the errors are obtained as the standard deviation from the mean value (See Materials and Methods). NB: no binding.

| Experiment | n | $K_b (M^{-1})$ | $K_d (\mu M)$ | $\Delta H$<br>kcal/mol | $-T\Delta S$<br>kcal/mol | $\Delta G$<br>kcal/mol |
| --- | --- | --- | --- | --- | --- | --- |
| 575-625 : RACK1 | 1 | $(3.7 \pm 0.4) \times 10^5$ | $2.7 \pm 0.3$ | $-5.7 \pm 0.1$ | $-1.9 \pm 0.2$ | $-7.6 \pm 0.1$ |
| 587-637 : RACK1 | 1 | $(4.4 \pm 0.3) \times 10^5$ | $2.3 \pm 0.2$ | $-7.8 \pm 0.1$ | $-0.2 \pm 0.1$ | $-7.7 \pm 0.1$ |
| 600-625 : RACK1 | 1 | $(3.9 \pm 0.6) \times 10^5$ | $2.5 \pm 0.4$ | $-5.2 \pm 0.2$ | $-2.4 \pm 0.1$ | $-7.6 \pm 0.1$ |
| 600-650 : RACK1 | 0.9 | $(4.7 \pm 0.5) \times 10^5$ | $2.1 \pm 0.3$ | $-7.2 \pm 0.2$ | $-0.5 \pm 0.2$ | $-7.7 \pm 0.1$ |
| 550-600 : RACK1 | NB | NB | NB | NB | NB | NB |
| 615-625 : RACK1 | 0.9 | $3.4 \pm 0.2$ | $3.0 \pm 0.1$ | $-8.1 \pm 0.1$ | $-0.6 \pm 0.1$ | $-7.5 \pm 0.1$ |
| 612-625 : RACK1 | 0.8 | $4.9 \pm 0.1$ | $2.0 \pm 0.1$ | $-8.3 \pm 0.1$ | $-0.6 \pm 0.1$ | $-7.7 \pm 0.1$ |
| 612-637 : RACK1 | 0.9 | $4.8 \pm 0.2$ | $2.1 \pm 0.1$ | $-8.6 \pm 0.1$ | $-0.8 \pm 0.1$ | $-7.8 \pm 0.1$ |

### CR2 comprises the LARP4 sequence that binds RACK1 propeller 5-7 region

A multiple sequence alignment (MSA) of fourteen sequences from varied vertebrates revealed conservation of the minimal RACK1 binding region that was nearly coincident with CR2 (32) (Supp Fig S2). Because a four amino acid insertion/deletion preceding this region distinguished the LARP4 and LARP4B, we expanded the analysis to the previously compiled fifty-eight CR2 sequences (32), displayed as sequence LOGOs in Fig 2A. This revealed differences independently conserved in LARP4/4A and LARP4B at CR2 positions 2, 5, and 10, and with greater apparent differential conservation at positions 14-15 (Fig 2A).

**Figure 2.**
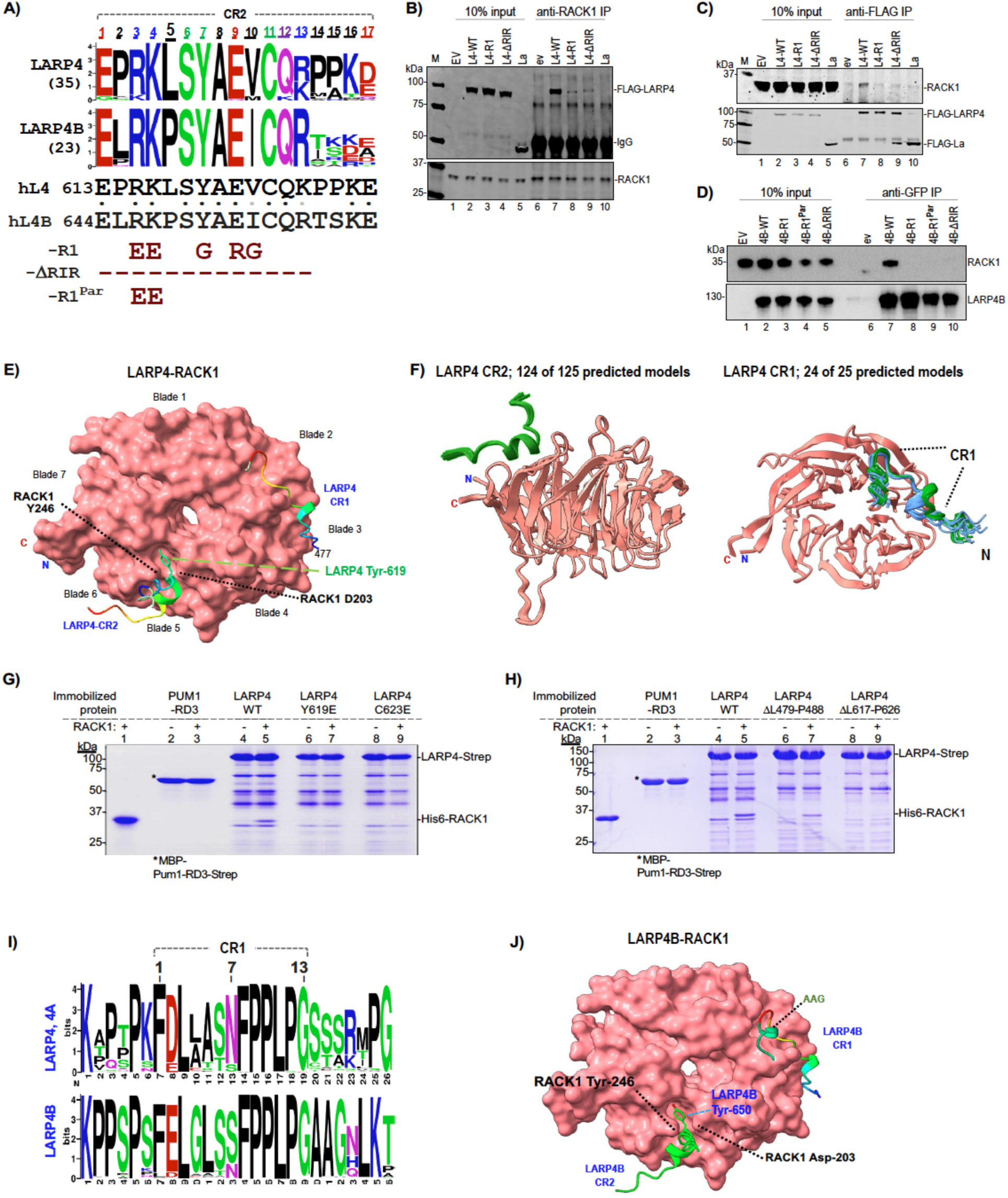
Mutations in conserved regions 1 and 2 of LARP4 and LARP4B disrupt association with RACK1. **A)** Top: sequence LOGOs (60) derived from the sequences in the CR2 MSA (32). Bottom: sequences of human LARP4 and LARP4B. Under this are indicated two types of mutations used for immunoprecipitation (IP) below; R1 has substitutions, ι1RIR has the positions deleted, and R1^Par^ has two substitutions. **B)** Immunoblot showing input extracts used for IP (lanes 1-5) and products of anti-RACK1 IP (lanes 6-10). Extracts were isolated from HEK293 cells transfected with the LARP4 (L4) constructs indicated above the lanes, EV = empty vector. The blot was incubated with anti-Flag (top) and anti-RACK1 Abs (bottom panel); the IgG heavy chain signal is indicated. M, size marker (LiCor). **C)** Immunoblot as in B) showing inputs (lanes 1-6) and products of anti-Flag-M2 IP (lanes 6-10). The blot was incubated with anti-RACK1 (top) and anti-Flag Abs (bottom). **D)** RACK1 and LARP4B immunoblots showing input extracts (lanes 1-5) and products of anti-GFP IP (lanes 6-10). Extracts were from PC3 cells transfected with the LARP4B constructs indicated above the lanes. Ab used on the blots are indicated to the right of the panels. **E)** AFMultimer predictions of RACK1-LARP4(401–724) interaction; sequence input for LARP4 was amino acids 401-724. Shown is the best model selected from five predicted models as per AFM (63) default mode on ColabFold (Supp Fig S3). RACK1 is shown in surface view. LARP4 residues were made invisible except for CR1 (477–489) and CR2 (613–629) colored rainbow, labelled as indicated. Orientation and annotation of RACK1 propeller blades is according to Nielsen (49). **F)** Results of AFM random seed iterative model predictions of RACK1 and CR2 and CR1 sequences. Left: 25 random seeds x5 models each using the 17-mer CR2, LARP4(613–629); sequence input was 613-629. The figure shows the converged predicted models (124 out of the 125 models). Right: 5 random seeds x5 models each using the 13-mer LARP4 CR1 within a larger sequence, LARP4(463–505); sequence input was 463-505. The figure shows the converged predicted models (24 out of the 25 models), which resolved into two clusters for its N-terminal helical turn (blue and green). **G)** Coomassie blue stained gel of *in vitro* pull-down binding results using recombinant StrepIItag-immobilized LARP4-WT or single substitution versions indicated above lanes 4-9, and immobilized MBP-Pum1-RD3-StrepII (68) which served as a control (lanes 2-3). Purified recombinant His6-RACK1 used as input for lanes 3, 5, 7 and 9 is shown in lane 1. After binding reactions and extensive washing were completed, the StrepIItag-immobilized protein and anything bound were eluted by biotin (lanes 2-9). **H)** Coomassie blue gel of *in vitro* pull-down results as in G) with immobilized LARP4-WT or the internal deletion versions indicated above lanes 4-9. **I)** Sequence LOGO of CR1 and flanking sequences for LARP4 and 4A (top) and for LARP4B (bottom) derived from sequences obtained from BLASTP search using nr-clustered database (NCBI) with an extended CR1 which produced ∼500 sequences with annotations LARP4, 4A and 4B from a variety of eukaryotes. The CR1 defined by Deragon is indicated by a bracket above and numbered. **J)** AFMultimer predictions of RACK1-LARPB(433–738) interaction; sequence input was amino acids 433-738. Shown is the best model selected from five predicted models per AFM (63) (Supp Fig S4A,D). RACK1 is shown in surface view. AAG points to a predicted helical turn that follows the CR1.

### Mutations to key CR2 residues of LARP4 and LARP4B impair interaction with RACK1

A mutant with five substitutions within the binding region was designated R1 and with the region deleted was designated ΔRIR (Fig 2A, bottom). Immunoprecipitation (IP) from HEK293 cell extracts expressing Flag-tagged LARP4-WT (L4-WT), LARP4-ΔRIR and LARP4-R1 was first performed with anti-RACK1 antibody (Ab). Fig 2B shows comparable expression of the Flag-proteins including negative control La (lanes 2-5, upper). Anti-RACK1 reproducibly IPed L4-WT (28) whereas L4-R1 and L4-ΔRIR were at lower levels as expected (lanes 6-10). Reciprocal IP using anti-FLAG followed by immunoblotting with anti-RACK1 Ab showed pulldown by L4-WT but not L4-ΔRIR, L4-R1 nor La, confirming intact CR2 is required for stable interaction (Fig 2C top).

LARP4B with corresponding R1 and ΔRIR mutations, plus a partial substitution mutant (R646E, K647E) named L4B-R1^Par^, and N-terminal GFP tags were used for anti-GFP-TRAP IP after transfection of PC3 cells (Fig 2D). L4B-WT co-IPed RACK1 whereas L4B-R1, L4B-R1^Par^ and L4B-ΔRIR did not (Fig 2D). The IPs validate Y2H and ITC data and show that the CR2 is critical for stable association of LARP4 and LARP4B with RACK1 in cells. In later sections we use the R1-CR2 mutant to examine effects on LARP4 activity. IPs were performed using different cells and routine methods in the different Laboratories in which they were done. Constructs other than these were not tested for RACK1 interaction in cells. Although it may appear from IP data that CR2 mutations have stronger effect on LARP4B than LARP4, there has not been systematic analysis.

### Protein-protein interface prediction algorithms elaborate LARP4-RACK1 interactions

Given recent developments in AlphaFold2-Multimer (AFM) algorithms to predict protein-protein interfaces (61,63), we obtained computational predictions of LARP4-RACK1 interactions. Although the predictions and associated experimental data were obtained after the results shown in later sections, their presentation is best in Figure 2. Using LARP4(401–724) and RACK1 as inputs, AFM predicted two LARP4 interaction sites, CR2, consistent with the binding data, as well as an additional site, spanning CR1 (Fig 2E). Convergence of multiple predicted structures was better at CR2 than at CR1 (Supp Fig S3A, video Fig S3B).

The high-resolution crystal structure of RACK1 alone reveals a hydrophobic cavity between its propeller blades 5 and 6, defined in part by the side chains of Tyr-246 and Asp-203 that project away towards the solvent (53). This cavity was modelled to be the LARP4 CR2 interface by AFM. To further examine the prediction of the CR2 site, the random seed iterative feature of AFM in ColabFold was used to increase sampling of the computational models (61). 124 of the 125 models using shorter sequences encompassing CR2 converged to a predicted helical turn spanning 619-YAEVC-623 in LARP4 CR2 located in this hydrophobic cavity of RACK1, strongly supporting the model (Fig 2F, left). The predicted interaction of CR2 with blades 5-6 is consistent with the Y2H data that identified RACK1 ORF codons 200-317 as the minimal interaction region (28) (Fig 1B).

The AFM models placing LARP4 619-YAEVC-623 in the crevice between blades 5-6 of RACK1 (Fig 2E,F) was validated by mutagenesis analysis using a protein-protein direct interaction assay (69,70). Immobilized recombinant FL-LARP4-Strep-tagII and mutants thereof were examined for interaction with purified recombinant FL-RACK1 (68). After coincubation, washing and elution with biotin, RACK1 was pulled down by LARP4 but not by the Pumilio RD3 domain negative control (68) (Fig 2G, lanes 2-5). The single substitution protein, FL-LARP4-Y619E largely impaired FL-LARP4 for RACK1 interaction (Fig 2G, lanes 6-7), as did FL-LARP4-C623E (lanes 8-9). These data demonstrate direct interaction between the two full length proteins, and that independent single substitutions of two highly conserved amino acids in the predicted LARP4 CR2 interaction site, Y619 and C623, largely impair binding. While no other CR2 substitutions were examined, a deletion (ΔL617-P626) also exhibited loss of binding (Fig 2H, lanes 8-9).

The CR1 in invertebrate sequences existed in LARP4 prior to gene duplication and CR2 (32). A majority of the invertebrate CR1 and LARP4B CR1 sequences have Ser at position-7, but not vertebrate LARP4/4A (32). LOGOs derived from MSAs representing CR1s from a wide variety of eukaryotes revealed lower overall conservation by LARP4/4A as compared to LARP4B (Fig 2I). Random seed AFM models revealed that 24 of 25 predicted models converged for the C-terminal part of LARP4 CR1 while the N-terminal predicted helical turn resolved into two clusters interacting with RACK1 propellers 2-3 (Fig 2F right, blue, green). The LARP4(Δ479-488) CR1 deletion reduced RACK1 pull-down relative to FL-LARP4 (Fig 2H, lanes 6-7).

We used AFM to generate computational predicted models of LARP4B-RACK1 interaction (Fig 2J). For CR2, the predicted LARP4B-RACK1 model was very similar to the LARP4-RACK1 model (Fig 2E, J). AFM default 5×1 models converged, supporting the CR1 and CR2 predictions (Supp Fig S4A-B). As with LARP4, 124 of 125 AFM models using the 17-mer CR2 of LARP4B converged to a predicted helical turn spanning YAEVC located in the hydrophobic cavity between propellers 5-6 of RACK1 (Supp Fig S4C). For CR1, positions 5-7 of LARP4 and LARP4B were predicted with high confidence as a helical turn while the invariant FPPLP was extended to blade 2; however, a second helical turn encompassing the conserved AAG sequence in LARP4B (Fig 2I) was predicted in ∼30% of the random seed models (Fig 2J, CR1 green, AAG).

### LARP4 activity for reporter mRNA stabilization; the assay system

The steady state levels of a mRNA accumulates after transfection of its plasmid is used to monitor stabilization by LARP4 (13,17,22). As reaching steady state requires multiple half-lives (71,72), the half-life (t½) of β-globin mRNA with a TNFα-ARE (β-glo-ARE) of ∼75 minutes assures this occurs well before harvest (73) (see 22). As the ARE accelerates mRNA decay by recruitment of CNOT (Introduction), the β-glo-ARE reporter focuses LARP4 activity to the deadenylation-linked decay pathway (15,22). LARP4 also increases levels of GFP mRNA and stable rp-mRNAs lacking ARE, on which it promotes accumulation of lengthened PATs (15,19,22) suggestive of the PAN2-PAN3 pathway (18). Thus, β-glo and GFP are controls for β-glo-ARE. The β-glo and β-glo-ARE reporters are distinguished only by the ARE (Supp Fig S5A and B). In northern blot assays, upward mobility shift of mRNAs results from LARP4-mediated PAT protection that manifests as increased PAT length that reflects slowed shortening during LARP4 expression after transfection (15,22).

Regarding the cotransfection system, data on mRNA poly(A) exists for LARP4 deletion as well as expression at endogenous levels (1X), 3-fold and 11-fold higher (15). Cells deleted of LARP4 exhibit transcriptome-wide median PAT length 5 nt shorter than WT, while cells expressing LARP4 3-fold higher than endogenous levels exhibit median PAT length 5 nt longer than endogenous levels (15). Thus, previous studies using the cotransfection approach provided evidence that it is a reliable assay system (15). Our standard method uses a L4-WT or L4-mutant expression plasmid added to equal aliquots of a mixture of the reporter plasmids to be compared in the same cells, typically yielding ≥90%, transfection efficiency.

### Mutations to CR2 decrease LARP4 activity to stabilize β-glo-ARE mRNA

L4-R1, L4-ΔRIR and other L4 constructs (Fig 3A) were examined for effects on three reporters, the stable mRNAs GFP and β-glo both lacking the ARE, and the β-glo-ARE-mRNA (54). Results typical from one of four replicate experiments are in Figure 3B-C; immunoblotting showed the expressed Flag-tagged L4 proteins (Fig 3B). A northern blot was probed for the reporter mRNAs and endogenous GAPDH as a loading control (Fig 3C). Quantifications of the three reporters normalized by endogenous GAPDH are shown in Fig 3D (upper graphs). In agreement with published data, L4-WT led to ∼2-fold higher levels of β-glo-ARE mRNA and ∼3-fold higher levels of GFP relative to EV (Fig 3D) (15,22). Although the lower mRNA levels in L4-ΔPAM2 vs. L4-WT cells lacked statistical significance, they share long-end mobility shifts that suggest shorter PATs (Fig 3C, lanes 2 and 3). This may reflect deficiency of L4-ΔPAM2 to compete with PAN2-PAN3-PAM2 for recruitment to mRNA PATs via PABP-MLLE (18).

**Figure 3.**
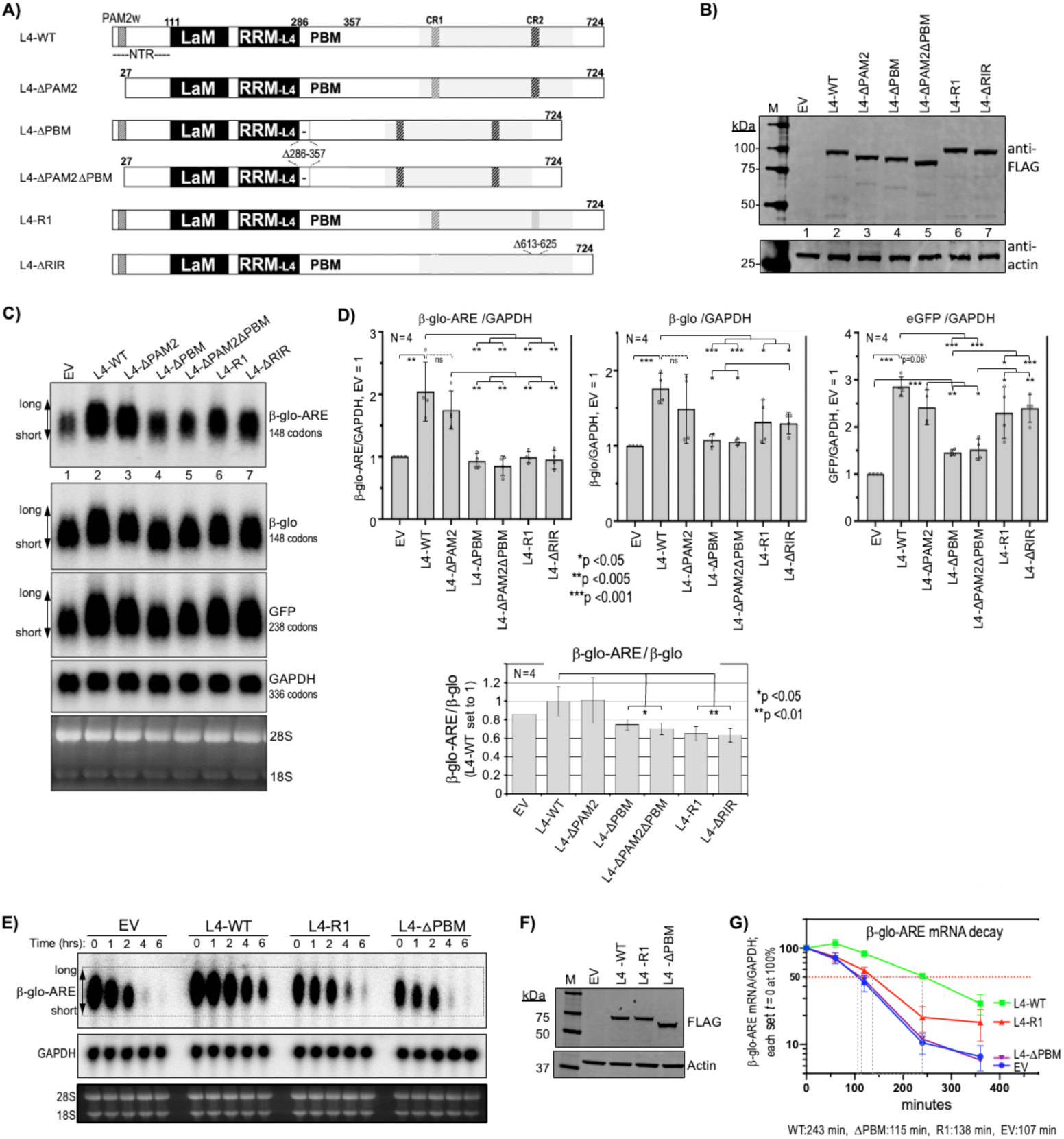
LARP4-RIR-CR2 mutants have lower ARE-mRNA stabilization activity than LARP4-ΔPAM2w. **A)** Schematic representation of the LARP4 (L4) constructs used in this experiment **B)** Immunoblot of extract protein from HEK293 cells transfected with the LARP4 (L4) constructs as in A) above the lanes; EV: empty vector, La: La protein (see text). **C)** Simultaneous responses of β-glo-ARE, β-Glo and GFP mRNAs in the same pool of transfected cells carrying a different LARP4 mutant. A northern blot of total RNA isolated from the same cells as in B probed for GFP, β-glo-ARE and GAPDH mRNAs as indicated in panels i-iv (labeled to the right); panel v shows a vertically compacted image of the EtBr-stained gel prior to transfer. **D)** Quantitation of northern blot signals of biological duplicate experiments. Transcript levels were normalized by GAPDH mRNA; N = 4; bar height represents the standard deviation (calculated using GraphPad Prism). **E-G) Analysis of β-glo-ARE mRNA decay.** HeLa Tet-Off cells were transfected with β-glo-ARE reporter and either empty vector (EV), LARP4-WT, LARP4-R1 or LARP4△PBM. Cells were harvested at 0, 1, 2, 4, and 6 hours after doxycycline addition to the media (Methods). **E)** Northern blot showing mRNA decay time course; samples and harvest times indicated above the lanes. **F)** Immunoblot of extracts isolated at time zero, processed for anti-Flag and anti-Actin antibodies as indicated. **G)** Graphic representation of quantified data of two β-glo-ARE mRNA decay experiments as in panel E. β-glo-ARE mRNA cpm was divided by GAPDH cpm in the same lane and the values at *t* = 0 for each of the four sets were set to 100% (Y-axis); N = 2; error bars represent standard deviation, SD (calculated with GraphPad Prism). Numbers under the graph represent apparent *t*½ (time at which 50% of *t* = 0 RNA remains).

Reproducibly, L4-ΔPBM and L4-ΔPAM2ΔPBM were most clearly impaired for the mRNAs (Fig 3D). L4-R1 and L4-ΔRIR were most impaired for β-glo-ARE mRNA, at p <0.005 relative to L4-WT and to L4-ΔPAM2 (Fig 3D, left). Although L4-R1 and L4-ΔRIR were significantly less active than L4-WT for β-glo mRNA, they were more active than L4-ΔPBM and L4-ΔPAM2ΔPBM (Fig 3D, middle). Further comparison revealed that while L4-R1 and L4-ΔRIR were as inactive as L4-ΔPBM and L4-ΔPAM2ΔPBM for β-glo-ARE stabilization, they each showed significantly greater activity for GFP (Fig 3D, right). Quantification confirmed L4-WT expression ∼3-fold higher than endogenous LARP4; however, even at 4-5-fold higher than endogenous LARP4, L4-R1 and L4-ΔRIR were critically deficient for β-glo-ARE stabilization but not GFP (not shown). Loss of the stabilizing effects of the ARE were best discerned by plotting β-glo-ARE/β-glo mRNA levels (Fig 3D, bottom). This showed that the LARP4-R1 and -ΔRIR mutants were impaired relative to LARP4-WT.

To examine effects of L4-R1 on β-glo-ARE stabilization, we assayed mRNA decay (22) (Fig 3E-G). β-glo-ARE mRNA levels were measured at time zero and at four time points following transcription inhibition in cells with L4-WT, L4-R1, and L4-ΔPBM expressed at similar levels (Fig 3F). This confirmed β-glo-ARE mRNA stabilization by L4-WT but not L4-ΔPBM (22), and showed L4-R1 was largely impaired (Fig 3G). Thus, CR2 mutations impair LARP4 activity for β-glo-ARE mRNA stabilization. That β-glo-ARE reveals this activity more than GFP and β-glo mRNAs likely reflects ARE-mediated CNOT recruitment and its shorter half-life. Accordingly, a larger percent of β-glo-ARE mRNA lifetime would be spent in the CNOT-decay pathway as compared to stable mRNAs engaged by PAN2-PAN3 unlinked to decay (42). These data might suggest that interaction with RACK1 would enhance LARP4 activity to oppose co-translational decay of β-glo-ARE mRNA.

### CR2 mutations impair LARP4 for stable association with ribosomes

As a conserved 40S-associated protein, RACK1 cosediments with translating ribosomes-polysomes in mammals, yeast and other eukaryotes (74,75). When RACK1 was purified to identify its mRNP-associated factors in brain, these included PABP and LARP4B (murine KIA0217) as prominent proteins (76).

Polysome profiles are used to compare translational status of mRNAs and related factors (55,58,77). The set of four HEK293 cell lysates were analyzed for LARP4 protein and reporter mRNA expression (Fig 4A-B) prior to examination by polysome profiles (Fig 4C). Figure 4A shows comparable expression of intact full length L4-WT and L4-R1. The L4-R1* is a control sample that differs from L4-R1 and the others by transfection with 2X β-glo-ARE plasmid so to produce comparable levels of β-glo-ARE mRNA in the L4-R1* and L4-WT cells. To address the possibility if 2X production of β-glo-ARE-mRNA would saturate the ARE-decay machinery (54), we note that artifactual stabilization would be expected to produce steady state levels greater than 2-fold higher than in controls (72). As shown to the right of the northern blot, quantification revealed ∼2-fold more β-glo-ARE-mRNA in L4-WT than in EV cells, whereas L4-R1 cells were inactive for increasing β-glo-ARE mRNA but active for increasing β-glo and GFP mRNA levels (Fig 4B). In the control L4-R1* cells, the β-glo-ARE mRNA expressed from 2X plasmid did not accumulate to greater than 2-fold higher than in L4-R1 cells (Fig 4B).

**Figure 4.**
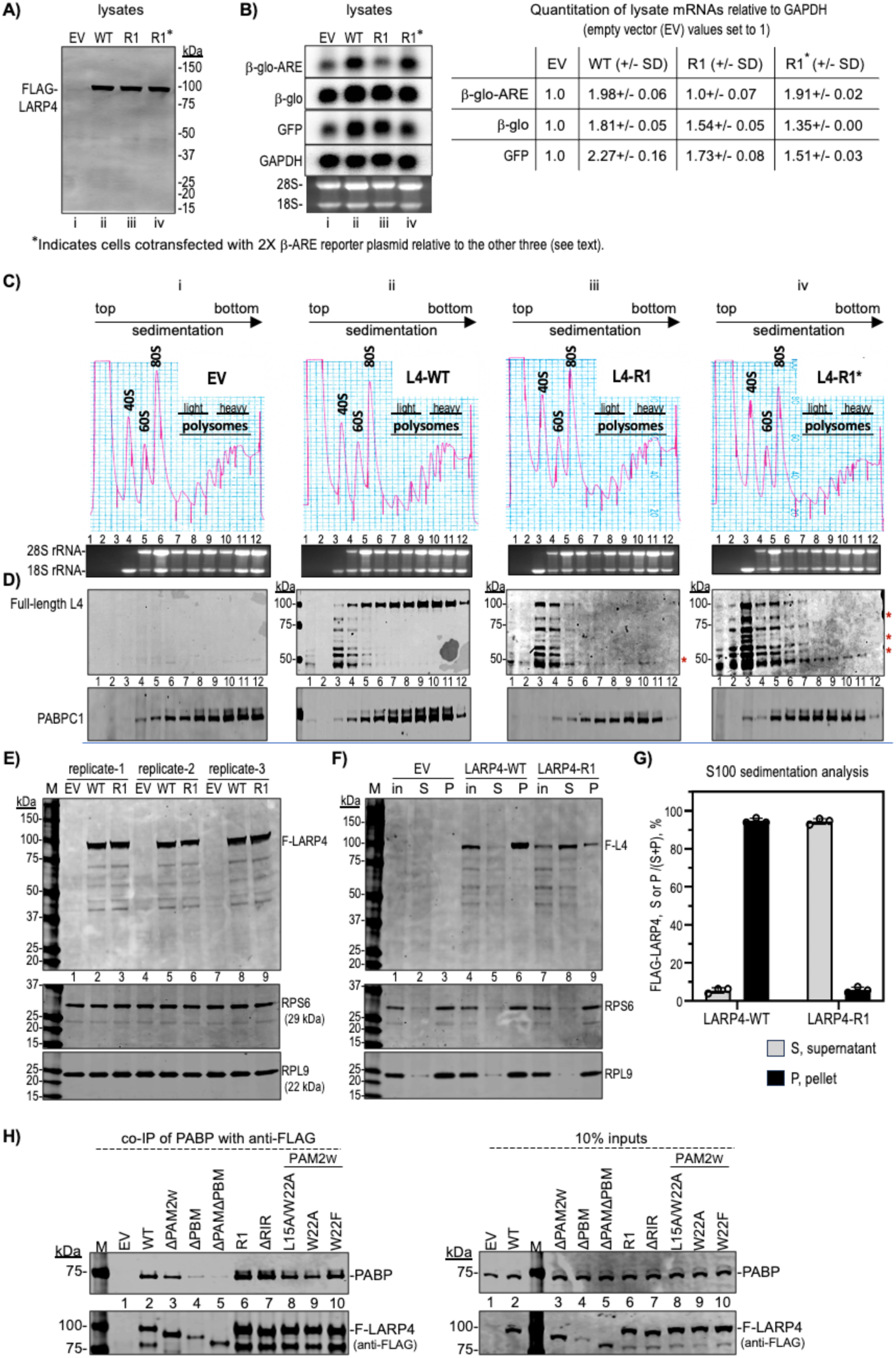
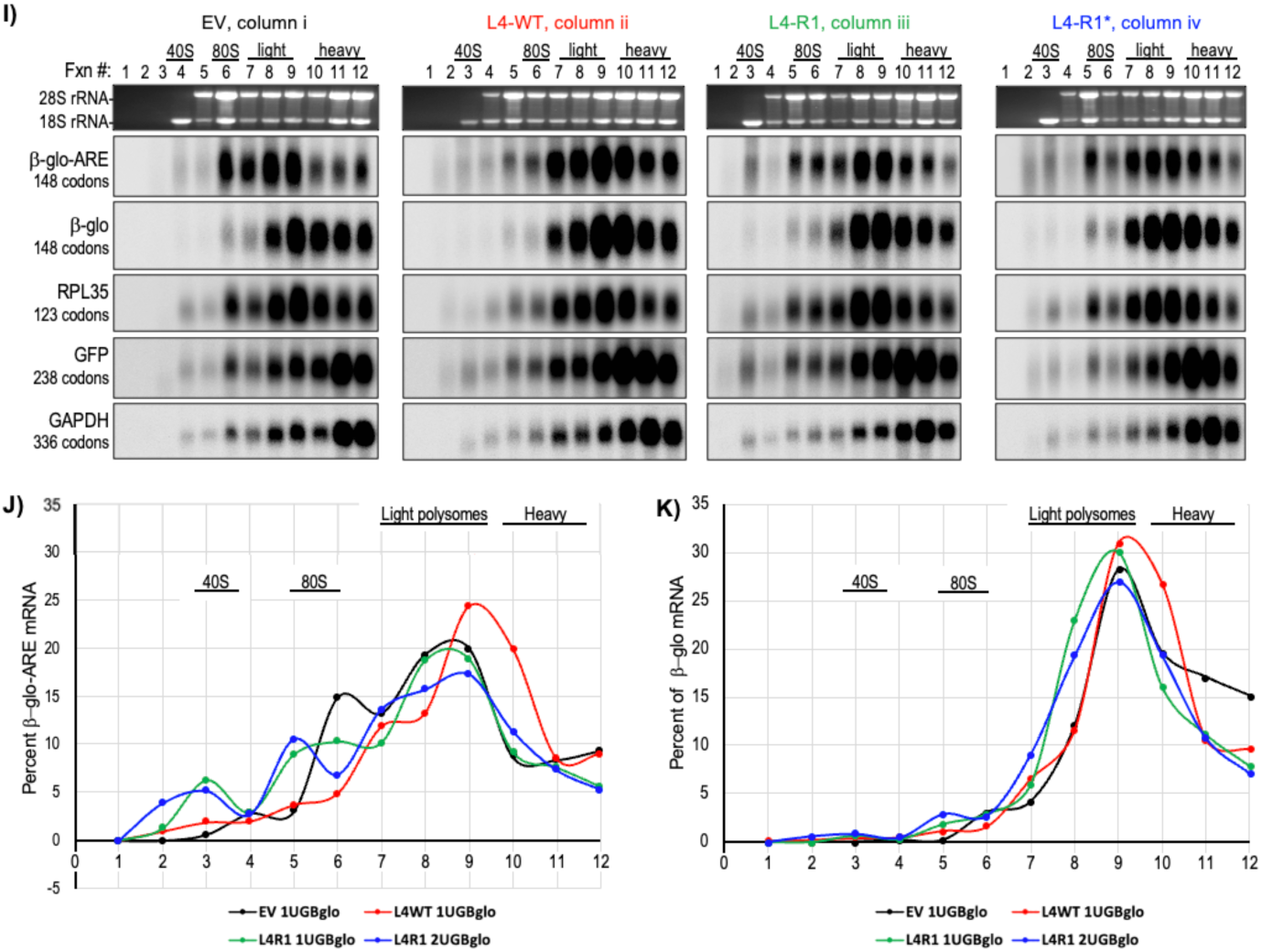
LARP4 exhibits CR2-dependent ribosome association and β-glo-ARE mRNA translation-related activity. **A-B)** Analysis of input cell lysates used for sucrose gradient sedimentation. **A)** Immunoblot of input lysates from samples with transfected constructs indicated above the lanes, EV (empty vector), L4-WT, L4-R1 and L4-R1* the latter with twice as much β-glo-ARE reporter as the others probed with anti-Flag Ab. **B)** Northern blot of RNAs from the same lysates, probed for mRNAs to the left with quantification of relative amounts shown to the right, based on replicates; N=4 for EV and WT, N =3 for R1 and N =2 for R1*. SD = standard deviation, calculated using Prism. Designations i, ii, iii, iv, under the lanes of A and B correspond to the four columns in C. **C)** Polysome profiles of the lysates in A-B run in parallel, EV, L4-WT, L4-R1 and L4-R1*; the 40S, 60S, 80S peaks and polysome regions are indicated. RNA and protein from each fraction (numbered) were analyzed. The EtBr-stained agarose gel shows 28S and 18S rRNAs under the profiles; note that 18S in absence of 28S is a marker for ∼40S ribosome subunits. **D)** Immunoblots with Abs to detect proteins indicated to the left; full-length LARP4 detected by anti-Flag Ab migrates just under the 100 kDa marker (upper panel). Molecular weight markers in kDa are indicated to the left (columns ii-iv); asterisks to the right of iii-iv indicate LARP4-R1 fragments. **E-G)** Sucrose cushion sedimentation S100 analysis. **E)** An immunoblot of three replicate lysates from EV, L4-WT and L4-R1 (equivalent amounts of OD260) analyzed for Flag-L4, RPS6 and RPL9 as indicated; lane numbers are provided under the upper gel blot. **F)** Representative immunoblot results of S100 sedimentation analysis showing the inputs (in), supernatants (S), and pellets (P) for the (replicate 1) lysates from EV, LARP4-WT and LARP4-R1 samples as indicated; lane numbers are under the upper blot. **G)** The Flag-LARP4 signals in the S and P fractions were quantified by LiCor and the fractions of each in the triplicates (Supp Fig S7) were plotted as S/(S+P) and P/(S+P), for LARP4-WT and for LARP4-R1, as indicated. **H)** Co-IP of PABP with anti-Flag for different Flag-LARP4 constructs. The Left panel shows products of the IPs, and the right shows the inputs; Δ denotes deletion constructs as in Fig 3A (see text). See Supp Fig S8A-C. **I)** Northern blot analysis of the mRNAs in the polysome fractions (C-D) as indicated to the left of the vertically stacked panels (codon ORF lengths are in parentheses). **J-K)** Quantification of the β-glo-ARE (J) and β-Glo mRNA (K) in the blots in each fraction expressed as percent of the combined total in all fractions (55,58).

Immunoblots using anti-FLAG Ab revealed full-length Flag-L4-WT enriched in 80S and polysome fractions, whereas apparent Flag-L4-WT degradation fragments were in the pre-80S fractions 3-5 (Fig 4D, column ii). By contrast, there was no enrichment of Flag-L4-R1 in 80S or polysome fractions, as the majority the Flag-L4-R1 signal accumulated in the pre-80S fractions mostly as fragments (column iii). Higher sensitivity detection revealed the same trend for the L4-R1* profile with evidence of Flag-containing fragments in polysome fxns (asterisks).

Similar results were reproducibly observed in additional independent experiments, one of which is shown in Supp Fig S6; less Flag-L4-R1 was on polysomes, and more was shifted to pre-80S fractions as compared to Flag-L4-WT. The apparent degradation occurred despite absence of protease use for detachment of cells from growth dishes prior to lysis, and the presence of protease-inhibitors in the lysis buffer. Thus, L4-R1 was intact in lysates, but susceptible to degradation associated with polysome profile preparation and analysis (Fig 4A vs. C). We therefore tested a less taxing method to examine ribosome cosedimentation.

Lysates from cells transfected with EV, L4-WT and L4-R1 were prepared as for polysome profiles, laid atop 1-ml sucrose cushions and subjected to ultracentrifugation for 1h so that ≥80S complexes pass through the cushion and form a pellet, and the rest remains as supernatant (57). Fig 4E shows an immunoblot of three sets of lysates probed for Flag-L4, RPS6 and RPL9 in the upper, middle, and lower panels respectively, the latter two representing ribosomal protein subunits. Fig 4F is one of the three immunoblots after sedimentation analysis showing the inputs (in), supernatants (S), and pellets (P). The majority of L4-WT sedimented with RPS6 and RPL9 in the pellet with little in the supernatant (Fig 4F, lane 6 vs. 5). By contrast, the majority of L4-R1 remained in the supernatant, while a minority sedimented with the pellet (lane 9 vs. 8). L4-R1 was not accompanied by RPS6 or RPL9 in the supernatant, which maintained sedimentation with pelleted ribosomes as expected (Fig 4F, lane 8 vs. 9). Triplicate analysis validated that L4-WT efficiently cosedimented with ribosome complexes while L4-R1 remained mostly in the supernatant as full-length protein (Supp Fig S7). Flag-L4 in S and P were quantified by LiCor and the fractions of each of the triplicates were plotted (Fig 4G).

In sum, LARP4-WT maintains stable association with RACK1 assayed by co-IP, and cosediments with ribosomes-polysomes. Substitution of key residues in CR2 disable LARP4-R1 to co-IP RACK1 and for association with ribosomes-polysomes in polysome profiles and by sucrose cushion sedimentation analysis.

### LARP4 RACK1-interaction mutants maintain interaction with PABP

RACK1 and PABP exhibited noncompetitive binding to a LARP4B fragment C-terminal to its RRM, consistent with PABP binding to the PBM region and RACK1 binding downstream (50) (Fig 1A). LARP4 mutations that impair interaction with PABP do not significantly alter distribution of the abundant PABP in polysome profiles (28). Here we examined LARP4-R1 and LARP4-ΔRIR for PABP co-IP, using multiple LARP4 PABP-interaction mutants to help gauge relative binding. As noted above, LARP4-PABP interaction involves the PBM and PAM2W, and the latter can be competed by poly(A) *in vitro* (31). Phe at PAM2 consensus position-10 and Leu at consensus-3 are the highest affinity determinants respectively for MLLE binding (18), represented by W22 and L15 in LARP4 (28). Diversity among twelve biochemically characterized PAM2 sequences (most were structurally characterized) is associated with ∼200-fold range in affinity to the MLLE of PABP (reviewed in 18) of which the PAM2W of LARP4 is in the lowest of the three affinity groups (18). Relevant is that LARP4-NTD proteins with PAM2W single mutations W22F, and W22A, bound MLLE *in vitro* while L15A did not (31). More distinctly, LARP4-NTD-W22F strongly impaired poly(A)-binding (31) (A.R. & R.J.M., unpublished).

Co-IP of PABP was performed with anti-Flag to pull down LARP4 proteins from HEK293 (Fig 4H); the L4 proteins in lanes 2, 6, 8-10 contain point substitutions and other lanes show deletions. Of the PAM2W mutants, W22F appeared least impaired whereas deletions ΔPAM2 and ΔPAM2ΔPBM were progressively more deficient for PABP co-IP by comparison to WT as expected (Fig 4H, left). Consistent with relative high dependence on PBM, L4-W22A and -L22A/W22A were mildly deficient for PABP co-IP. Importantly, the RACK1 interaction mutants, L4-R1 and L4-ΔRIR appeared least impaired (Fig 4H, left, lanes 6-9). These results are corroborated by three independent co-IP data sets in Supp Fig S8, are in agreement with noncompetitive binding by RACK1 and PABP to LARP4B (50), and the notion of CR2, PAM2w, and PBM as separate motifs through which LARP4 can modulate mRNA-related activities.

### Polysome profiles suggest LARP4 promotes translation of β-glo-ARE mRNA

mRNAs in pre-initiation and pre-elongation complexes generally sediment in 40S-80S fractions, while translating mRNAs sediment deeper, although generally limited by codon ORF length and thus capacity for ribosome occupancy (55). The variability with which an mRNA sediments beyond 80S reflects its ribosome density and translational efficiency (TE). A northern blot with RNA from each profile in Fig 4C was probed for the mRNAs noted to the left of Fig 4I. The EV profile blot suggests lower TE for β-glo-ARE mRNA than for β-glo mRNA in the panel below it (Fig 4I, column i). Thus, differential sedimentation reflects that β-glo mRNA exhibits higher TE than β-glo-ARE mRNA, which differ only by the ARE, in the same cells (also see Supp Fig S6). These results are consistent with other data indicating that in addition to mediating mRNA decay, the TNFα ARE can impose negative effects on translation via TTP-mediated recruitment of repressor proteins (41,78–83).

Extending the analysis, the β-glo-ARE mRNA exhibited higher TE in L4-WT cells than in EV cells. In L4-WT cells, β-glo-ARE was more like β-Glo and RPL35 mRNAs (Fig 4I compare columns i, ii). Reversal of inefficient translation of β-glo-ARE mRNA in L4-WT cells is evidence against an unforeseen defect in the β-glo-ARE construct. Thus, TE of β-glo-ARE mRNA was increased by LARP4. This is a novel illustration of a distinct LARP4 activity, to promote translation of β-glo-ARE mRNA.

### LARP4-R1 is deficient for promoting cosedimentation of β-glo-ARE mRNA with polysomes

L4-R1 cells showed more left-shift of β-glo-ARE mRNA peak distribution and fractional distribution among light and heavy polysomes as compared to L4-WT (Fig 4I, columns ii and iii); this shift is observed in the percent of mRNA in each fraction, as plotted in Fig 4J. Notably, twice as much β-glo-ARE mRNA in fraction 10 in L4-WT cells relative to EV cells and L4-R1 cells, would represent higher potential TE for this mRNA (84). These data are evidence that L4-R1 is less active than L4-WT for promoting TE (Fig 4I, J). Although it seemed unlikely that lower levels of β-glo-ARE mRNA due to impaired mRNA stabilization by L4-R1 might account for its lower activity to promote TE, we designed a control experiment to examine this. L4-R1* cells express the same amount of β-glo-ARE as in L4-WT (Fig 4B) and found a left-shift like L4-R1, thus the impaired TE activity of LARP4-R1 was not rescued by increasing the levels of the β-glo-ARE (Fig 4I, ii-iv).

Because splitting of peak fractions occurs asymmetrically in some of the profiles relative to replicates, quantification of our triplicate profiles for amount of β-glo-ARE mRNA was by combining fractions representing light and heavy polysomes, or pre-polysomal fractions as percentages of total (see Figure 8-Supplement 4 in (55)). This showed that LARP4-WT promotes more β-glo-ARE mRNA on polysomes and less in pre-polysomal fractions with higher ratio and higher statistical significance (p <0.001) than either LARP4-R1 or EV (Supp Fig S6M). The triplicate sample profile analysis of β-glo-ARE mRNA in Supp Fig S6N shows that LARP4-R1 and EV are deficient relative to L4-WT to clear fractions 1-3 and 4-5 (including 40S-80S pre-polysomal translation complexes) and efficiently occupy polysome fractions 8-10.

We note that the differences observed between L4-WT and L4-R1 in polysome sedimentation profiles are manifest most robustly by β-glo-ARE mRNA. However, the L4-WT activity extends to β-glo mRNA which occupies heavy polysomes more efficiently in LARP4-ET cells than in LARP4-R1 cells (Fig 4K). This is reminiscent of differential effects on mRNA stability (Fig 3), namely that the effects are manifest more on the mRNA containing the ARE than on the mRNAs lacking the ARE.

### LARP4 PAT protection activity can increase translation efficiency of nanoLuc mRNA

While polysome profiles provide information about mRNA association with pre-initiation complexes and ribosome density, assessing efficiency of mRNA translation into polypeptide products requires additional assays (84). For this, mRNA reporters with destabilization elements that encode proteins with destabilization elements have been useful (85,86). The engineered NanoLuc luciferase (nLuc) and its novel assay reagent comprise a high activity reporter system even if the enzyme is appended with a C-terminal PEST protein degradation motif (nLucP) that decreases its half-life to 20 minutes (87). The TNFα-ARE added to the 3’ UTR of nLuc has been used to examine response to TTP and the innate inflammatory response in HEK293 cells (88) (see 89). We put the TNFα-ARE in the nLucP construct 3’ UTR. Similar to a published study (85), the PEST element and the ARE independently decreased nLuciferase activity, and combining them led to further decrease (below). After characterizing expression from our transfected nLuc reporters, we stably integrated them into a targeted chromosomal site in Flp-In HEK293 cells.

We devised an experiment that examines if increasing amounts of LARP4-mediated PAT protection would increase translation of nLucP-ARE mRNA, measured as luciferase activity, in conditions in which the nLucP-ARE mRNA levels per se were not significantly increased. The nLucP-ARE reporter plasmid was cotransfected with increasing amounts of LARP4-WT expression plasmid. Cells were harvested after 48h, of which 50% was used to quantify LARP4 levels and luciferase activity, and 50% was used for RNA quantification. LARP4-WT protein levels increased in a plasmid-dose dependent manner (Fig 5A-C). This was accompanied by upward gel mobility shift of nLucP-ARE mRNA characteristic of increased PAT length due to slowed deadenylation (15,22) while the mRNA levels varied relatively little (indicated under the lanes; Fig 5D). Thus, effects of LARP4-mediated PAT protection could be examined in conditions not confounded by increase in the mRNA levels (Fig 5D). The upper graph in Fig 5E shows nLucP luciferase activity/μg extract versus LARP4-WT protein levels, and the lower graph shows TE, (nLucP luciferase activity/μg extract)/(nLucP-ARE mRNA/μg extract) on the Y-axis versus LARP4-WT levels (X-axis), A >4-fold increase in translation of nLucP-ARE mRNA reflected by luciferase activity accompanied the dose-dependent LARP4-mediated PAT protection while levels of nLucP-ARE mRNA changed relatively much less. For this experiment, the R^2^ trend line derived from six data points with R^2^=0.93 provides evidence that LARP4 promotes TE of nLucP-ARE mRNA as monitored for functional enzyme product, complementing the conventional polysome profile fractionation (Fig 5E). This effect was independently reproduced using a partial codon-optimized version of LARP4 referred to as LARP4-CS (codon swap) (22), which yielded similar results with R^2^ >0.96 for TE (Supp Fig S9). These data are evidence that LARP4-mediated poly(A)-end protection (preservation of PAT length) can increase mRNA translational efficiency by means other than increasing the mRNA levels per se.

**Figure 5.**
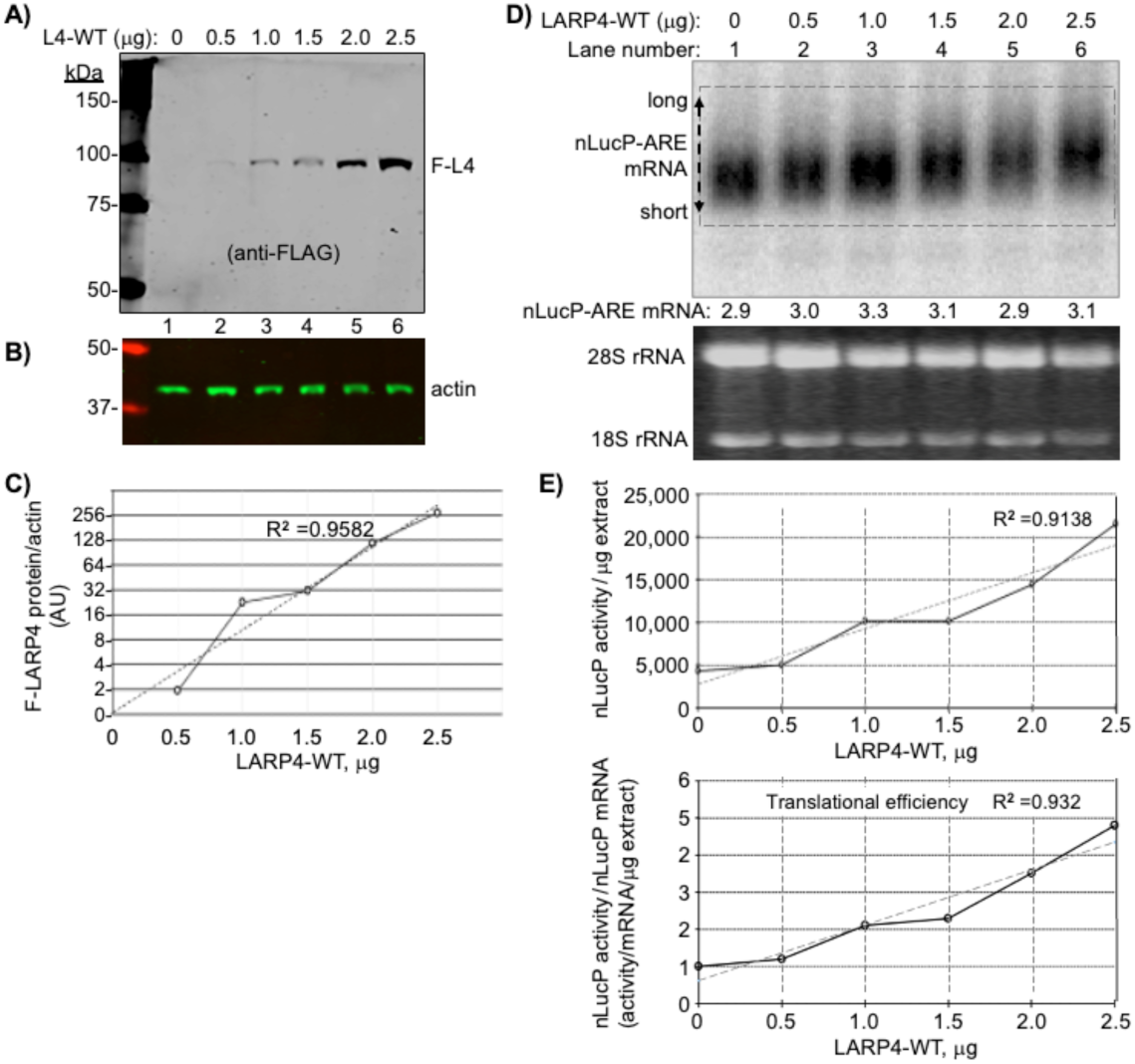
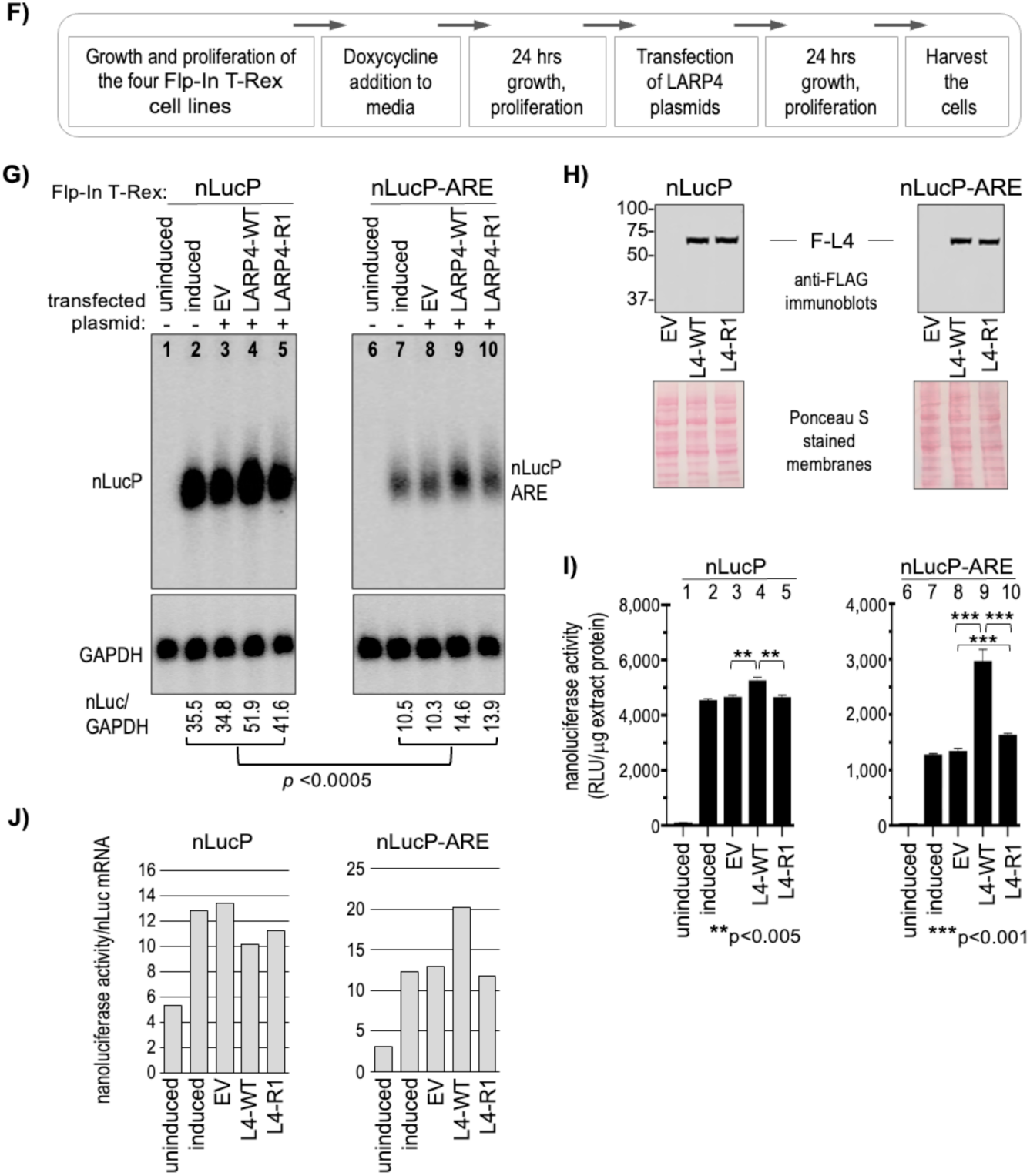
LARP4 activity increases translation efficiency of nanoLuciferase (nLuc) reporter mRNA. **A-E)** Analysis after transient transfection of nLucP-ARE reporter and LARP4 expression plasmids. Forty-eight hours after standard cotransfection of a nLucP-ARE and LARP4-WT expression plasmids (both under control of CMV promoter) cells were harvested and half processed for protein and half for RNA. **A-C:** Immunoblot analysis of LARP4-WT expression. To each of six transfection reaction mixtures containing equal amounts of the nLucP-ARE reporter plasmid, was added the different amounts of LARP4-WT plasmid in μg indicated above the lanes (0 = empty plasmid). The blot was developed using anti-Flag (A), anti-actin (B), and the results were quantified and graphed (C). **D:** Northern blot analysis of nLucP-ARE mRNA expression in the same samples as in A-B as indicated above the lanes. The nLucP-ARE RNA reactive with the probe was quantified and is indicated below the lanes. The double ended arrow at the left annotated with long and short refers to poly(A) length differences characteristic of LARP4 activity (see text). A dashed line rectangle helps discern the lower and upper boundaries of that reflect poly(A) lengths that contribute to the mobility shifts (13,15,90). **E:** The upper graph plots nLucP luciferase activity in relative light units (RLU)/μg extract (Y-axis) versus LARP4-WT protein levels (X-axis). The R**^2^** value derived from a linear fit trendline shown as a dashed line. The lower graph plots TE, (nLucP luciferase activity/μg extract)/(nLucP-ARE mRNA/μg extract) on the Y-axis versus LARP4-WT levels (X-axis). **F-J)** Analysis of two nLuc reporters integrated in Flp-In HEK293 cells, and responses to LARP4-WT and LARP4-R1. **F:** Diagram showing experimental schema. **G:** Northern blot analysis. The two panels contain RNAs from cells treated as indicated above the lanes, from the Flp-In T-Rex lines containing the single copy reporter: nLucP and nLucP-ARE. All samples were processed side-by-side; the two sets of samples were run on different gels and transferred to different blots which were then hybridized with the same antisense nLuc probe, washed together, and exposed on the same phosphorImager screen. Quantification of the nLuc/GAPDH RNA signals for each lane is shown below the panels. **H:** Immunoblot analysis of the Flag-LARP4 levels as indicated. **I:** Nanoluciferase activities in the protein extracts corresponding to the samples in G; note different scales on Y-axes. **J:** Quantitation of translation efficiency (TE, (nanoluciferase activity/nLuc mRNA) derived from data in I and mRNA levels.

### LARP4 activity to promote TE of nLucP-ARE mRNA is impaired by deficiency of RACK1-interaction

To eliminate variation that might be associated with cotransfection of the reporters, we created FRT-targeted integrants in Flp-In™ T-Rex™-293 cells. Clones bearing a single copy of one of four reporters, nLuc, nLucP, nLuc-ARE and nLucP-ARE, were grown, induced for transcription, transfected side-by-side according to Fig 5F, and processed for expression of nLuc mRNA, F-LARP4 and luciferase activity. Analysis of the four samples revealed >200-fold difference in basal luciferase levels between the highest and lowest activity reporters, i.e., with and without the PEST sequence, nLuc and nLucP-ARE, respectively (Supp Fig S9I). The data are consistent with >4 day half-life for the nLuc enzyme (lacking PEST) (87), and suggest >2 weeks to achieve steady state after induction of expression, vs. <2 hrs. for nLucP. Thus, high activities of constructs lacking PEST and the pre-steady state accumulation conditions that diminish reliability of measured effects of LARP4-WT vs. LARP4-R1 suggest we limit analysis to the nLuc-PEST constructs, nLucP and nLucP-ARE (Fig 5G-I).

The northern blot lanes of each set compare uninduced and induced cells, and the next three compare EV, expression of L4-WT and L4-R1. Immunoblot levels of FLAG-LARP4 are in Fig 5H. The nLucP-ARE led to significantly lower mRNA levels compared to nLucP (Fig 5G). The difference in luciferase activities in the EV samples, between nLucP and nLucP-ARE reflected the difference in their nLuc mRNA levels, ∼3.5-fold, reproducible in the induced samples (Fig 5I, lanes 2-3 and 7-8, compared to corresponding lanes in Fig 5G).

Nanoluciferase activity levels produced from nLucP-mRNA were higher in L4-WT than in L4-R1 cells and in EV cells by ∼15% (Fig 5I), whereas activity levels from nLucP-ARE mRNA were ∼200% higher in L4-WT cells than in L4-R1 and EV cells (Fig 5I). The larger effect of LARP4-WT on ARE-containing vs. ARE-lacking mRNAs is reminiscent of the data for β-glo-ARE. We next focused on nLucP-ARE mRNA as a substrate of L4-WT and L4-R1. The effects on TE and specificity were apparent when luciferase activity was normalized for nLuc mRNA levels as shown in Fig 5J. LARP4-WT robustly promoted TE of nLucP-ARE mRNA while LARP4-R1 was deficient for this activity on nLucP-ARE mRNA. While LARP4-WT led to increased luciferase activity relative to EV and LARP4-R1 from LucP lacking the ARE (Fig 5I), this reflects stabilization that led to more nLucP mRNA, though to high levels not well coupled to translation efficacy in this system (Fig 5J). Whereas by contrast, LARP4-WT effects on nLucP-ARE were to increase both stabilization and TE while LARP4-R1 was impaired (Fig 5J) (Discussion). We next further compared nLucP-ARE mRNA as a substrate of L4-WT and L4-R1.

### Polysome profile data support LARP4 CR2-mediated efficient translation of nLucP-ARE mRNA

The Flp-in nLucP-ARE cells were induced, transfected with EV, L4-WT, L4-R1, grown to comparable confluency, processed and lysates prepared. Prior to sedimentation, the lysates were examined for expression of nLucP-ARE mRNA, Flag-LARP4, and nanoluciferase activity/μg extract (Fig 6A-C). After normalization, the data showed higher translation efficiency of nLucP-ARE mRNA in L4-WT than in EV cells, and that the CR2 mutations impaired this activity in L4-R1 cells (Fig 6D), reproducing the results for nLucP-ARE in Fig 5G-J.

**Figure 6.**
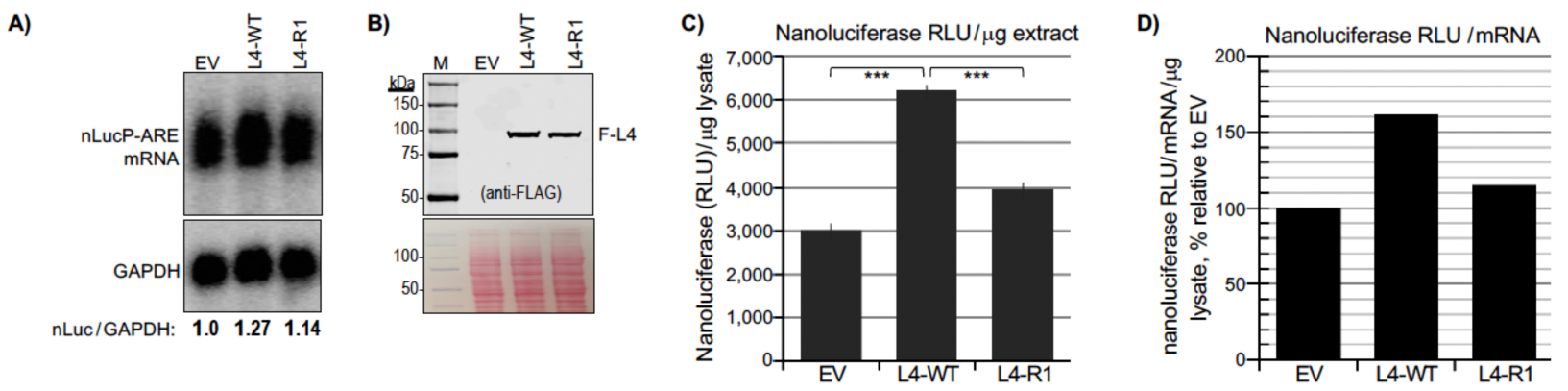
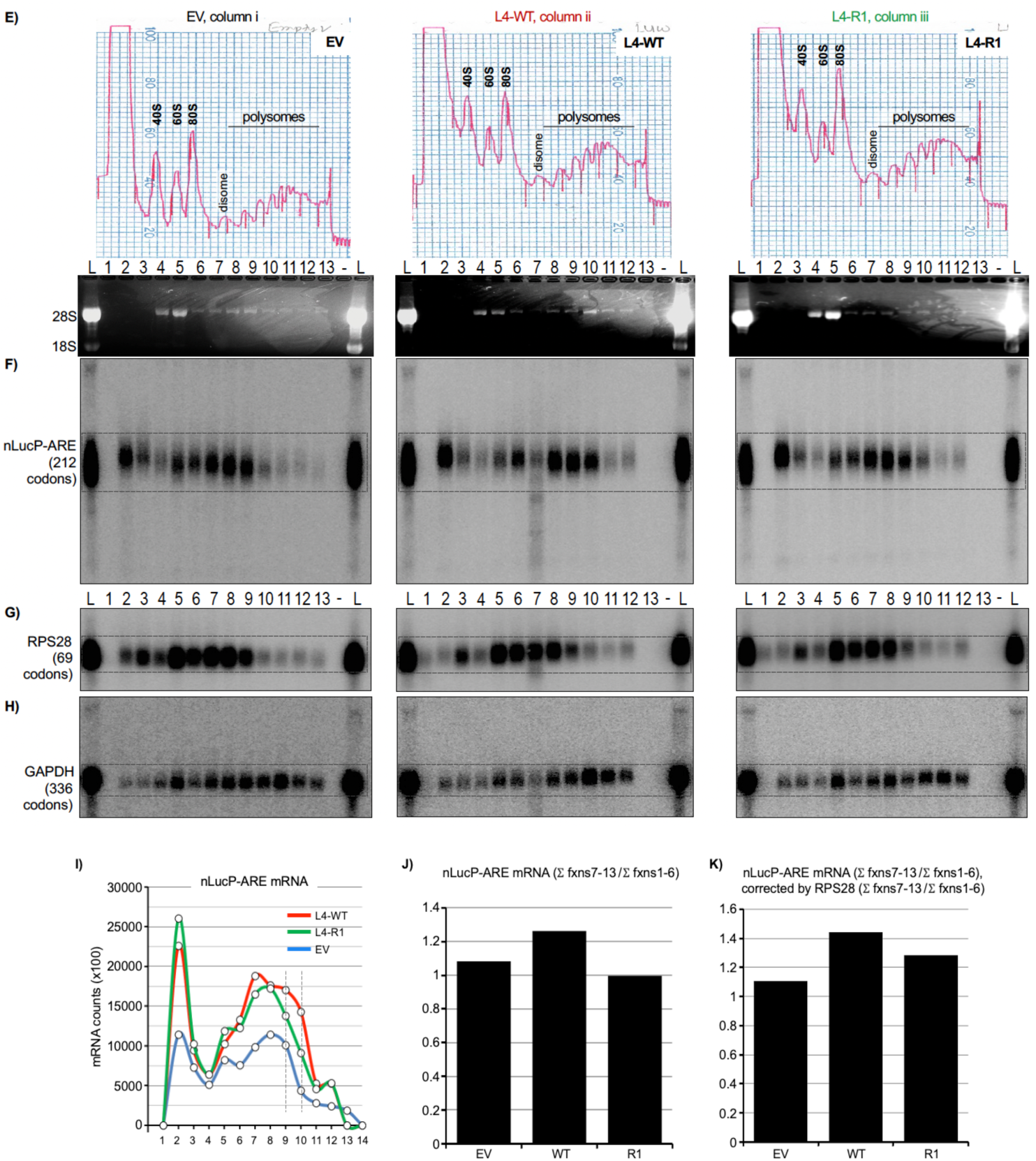
LARP4 activity to promote TE of nLucP-ARE mRNA is supported by polysome analysis, confirming β-glo-ARE analysis and dependence on the CR2 RACK1-interaction motif. **A-D)** Analysis of luciferase activity in Flp-In HEK293 nLucP-ARE cell lysates prepared for polysome profile sedimentation; cells had been transfected with EV, L4-WT and L4-R1, as indicated in the figures. **A:** Northern blot; Quantification of the nLuc signal/GAPDH is shown below the lanes, normalized to EV = 1.0. **B:** Immunoblot using anti-Flag Ab to detect Flag-LARP4 in the lysate. **C:** Graph of amounts of luciferase activity in RLU (relative light units) per μg lysate corresponding to the samples in A and B. **D:** Graph of amounts of luciferase activity in RLU per μg lysate, corrected for nLucP-ARE mRNA in each lysate, corresponding to the samples in A and B. **E-K)** Sucrose gradient sedimentation, polysome profile analysis of the three Flp-In HEK293 nLucP-ARE cell lysates represented in A-D. **E:** Columns i, ii, and iii show the OD254 nm tracings under which are the numbered fractions and Ethidium Bromide-stained agarose gel with 28S and 18S rRNA markers representing the 40S and 60S ribosome subunits. **F-H:** A northern blot from each polysome profile fractionation was probed for the three mRNAs indicated to the left, nLucP-ARE (F), RPS28 (G), and GAPDH (H) (codon ORF lengths are in parentheses). **I-K)** Graphic representations of nLucP-ARE mRNA distribution among polysome and pre-polysome fractions in EV, LARP4-WT and LARP4-R1 cell samples. **I:** Distribution of nLucP-ARE mRNA counts in each fraction of the polysome profiles as indicated. The vertical lines are to illustrate differences among the samples in the heavy polysome fractions 9 and 10 (see text). **J:** Bar graphs showing the sum of nLucP-ARE mRNA counts in polysome fractions 7-13 divided by the sum of nLucP-ARE mRNA counts in pre-polysome fractions 1-6. **K:** the same as in J but corrected by the amounts of RPS28 mRNA in the same profiles.

The lysates were subjected to sucrose gradient sedimentation. Tracings of the profiles upon collection are shown in Fig 6E columns i-iii. A northern blot from each profile was probed for mRNAs indicated to the left of Figs 6F-H (codon number in parentheses); in addition to the fractions, aliquots of total RNA from the lysate (L) were loaded as markers. In the EV profile (column i), most nLucP-ARE mRNA sedimented similarly to RPS28 whose relatively shorter codon number and ribosome occupancy limits it to fractions 8-9 with substantially less in fraction 10, whereas GAPDH sedimented deeper (Fig 6F-H, column i). By contrast, in lysate from L4-WT cells, the nLucP-ARE mRNA sedimented deeper than in EV cell extract, with a substantial amount in fraction 10, and deeper than the internal control, RPS28 in the same gradient (column ii) (although degradation occurred in fraction 7, this was incomplete and the RNA above background could be quantified, see below). Thus, the effect of LARP4 to promote the apparent TE on nLuc-ARE mRNA here are reminiscent of its effects on β-glo-ARE mRNA, and generally confirm this activity. The L4-R1 profile was deficient in shifting the distribution of nLucP-ARE mRNA relative to LARP4-WT. Most notably, the amount of nLucP-ARE mRNA in lane 10 in LARP4-R1 cells was decreased relative to lane 10 in L4-WT cells compared to the RPS28 pattern. Fraction 10 is especially relevant because in it reproducibly contains RNA from two polysome peaks (Fig 6E, also see Fig 4C and Supp Fig S6B) and therefore represents a larger increase in ribosome density relative to the preceding fractions, than the other fractions. The nLucP-ARE mRNA in each fraction is plotted in Fig 6I.

Fig 6J shows the sums of mRNA counts in fractions 7-13 divided by the sums in fractions 1-6, i.e., polysomes/pre-polysomes (55) and Fig 6K shows this after normalization by RPS28 mRNA. The data in Figs 6I-K indicate a larger amount of the nLucP-ARE mRNA on polysomes in LARP4-WT cells than in LARP4-R1 or in EV cells. While these data do not account for disproportionately large effects on TE of two polysome peaks in fraction 10, they are consistent with and support the more functional output assessment of TE based on luciferase activity measurements in Fig 6D.

## DISCUSSION

A major conclusion from this work is that the short conserved region (CR2) of LARP4 (amino acids 615-625) comprises the principal site that interacts directly with RACK1. Substitution and/or deletion mutations to CR2 and the homologous region of LARP4B markedly impair stable interaction with cellular RACK1. These mutations strongly impair stable association of LARP4 with ribosomes and polysomes relative to intact LARP4. Use of model reporters showed that the mutations decrease LARP4 activity for stabilization of β-glo mRNA containing an ARE to significantly greater extent than stabilization of the stable β-glo mRNA lacking the ARE.

Dissociation from polysomes of RACK1 interaction-deficient LARP4-R1 protein was accompanied by its susceptibility to proteolysis in the several sucrose gradient sedimentation experiments performed. That the LARP4-R1 proteolysis could be minimized by use of milder cosedimentation conditions indicated that polysome dissociation occurred independent of proteolysis. The CR2 mutations (R1 and ΔRIR) which caused loss of LARP4-RACK1 interaction by IP assay caused minimal if any loss of LARP4-PABP interaction by IP assay, in agreement with published data on LARP4B (50). The data would suggest that RACK1 and PABP can independently modulate LARP4 mRNA-related activity via their separate binding sites on LARP4. A caveat is that the IP assays do not probe functional relationships among these proteins. The LARP4-R1-specific proteolysis during polysome profile analysis confounded our ability to examine of this aspect of function.

Another significant conclusion is that LARP4-WT was shown to robustly promote the TE of both β-glo-ARE and nLucP-ARE. Further, the CR2-RACK1 interaction motif of LARP4 was shown to promote TE of these ARE-containing reporter mRNAs. This shouldn’t be surprising as the CR2 motif physically links LARP4 to the ribosome via direct binding. Evidence of this were conventional polysome profiles which display ribosome density, that compared β-glo-ARE and β-glo mRNAs. This was then confirmed using nLucP and nLucP-ARE mRNAs which allow characterization of TE in terms of translation output per unit of mRNA as functional luciferase enzyme activity.

LARP4 activity dependence on the CR2 was manifested most robustly for the β-glo-ARE and nLucP-ARE mRNAs relative to their controls lacking the ARE, it was also manifest albeit less so for the more stable controls which lack the ARE. The CR2-dependent LARP4 activity for nLucP-ARE mRNA was examined for TE in the same extracts used for polysome profile analysis (Fig 6). The polysome profiles revealed that while LARP4-WT promoted high ribosome density on nLucP-ARE mRNA, the LARP4-R1 (CR2) mutant was deficient by comparisons to internally controlled RPS28 mRNA in the same cells on the same profiles (Fig 6).

Yet while polysome profile analysis of nLucP-ARE mRNA confirm the data obtained using β-glo-ARE, and support the TE data based on translation output, luciferase activity/nLucP-ARE mRNA (Fig 6C-D), we noted a feature of the nLucP-ARE mRNA profiles that reproducibly and intriguingly differed from the β-glo-ARE profiles, substantial amounts of nLuc-ARE mRNA in fractions 2 (Fig 6E-H and I). The material in fraction 2 represents soluble mRNP not engaged by 40S components, at levels higher than in any other fraction thereby comprising sizable percentages of the nLucP-mRNA which was not observed for the other mRNAs in Fig 6 nor for β-glo-ARE mRNA. Although other potential explanations exist, the possibility that this is related to the PEST sequence is considerable and raises questions for future studies. As shown in Supp Fig S9G, we found that the PEST sequence reproducibly conferred significant stabilization to nLuc mRNAs independent of whether or not they contained the ARE. It is thus tempting to suggest that this PEST sequence may account for accumulation of nLucP-ARE mRNA in the fractions 2. If so, some PEST sequences might act as an RNA stabilization element in the non-translating mRNA, and a destabilizing element in its translated product.

The mRNA metabolism and translation related work here were done using cotransfection of HEK293 cells that carry endogenous LARP4. Previous studies compared LARP4 knockout (KO) and WT cells for effects on endogenous mRNA stability and transcriptome-wide PAT metabolism, and included HEK293 cells cotranfected with mRNA reporters and plasmids expressing LARP4-WT at various levels (15). Although those studies produced evidence that the LARP4 expression-in HEK293 LARP4-replete cell system would reflect meaningful results (15), we created HEK293-derived LARP4-knockout (KO) cells by CRISPR-Cas9 and transfected them with LARP4-WT and LARP4-R1 to examine β-glo-ARE mRNA metabolism (Supp Fig S11). While there is no a priori reason to believe that such cells have not acquired compensatory mechanisms and are better than the replete system, the data obtained nonetheless readily supported the basic results as expected and confirm them. Specifically, LARP4-WT and LARP4-R1 exhibited differential activity for stabilization of β-glo-ARE mRNA vs. the GFP mRNA lacking the ARE, reproducibly, and with statistical significance (Supp Fig S11).

### Context of the RACK1 interaction region

LARP4 and LARP4B exhibit similar overall architectures but with different functionalities in cancer and other processes (33), verified by direct comparisons (37). Compared to their N-terminal halves which share multiple distinct motifs, their second halves display no recognizable domains and relatively low sequence identity with each other except at CR1 and CR2 (Fig 1A). The work here revealed the significance of CR2, and possibly CR1. As suggested by data in figure 2, cell type differences and differences between LARP4 and LARP4B in their interactions with RACK1 may exist, though such possibilities have not been examined.

We wondered if the predicted LARP4 binding sites on RACK1 would appear accessible in context of the ribosome. Inspection of cryo-EM structures of human 80S ribosomes (PDB 6Z6M (91) (92)) suggested the predicted CR2 and CR1 interaction sites as accessible, located on the RACK1 side facing outward (**Supp Fig S10A**, upper and lower). AFM predictions were performed using RACK1, C-terminal region of ribosomal protein uS3 which interacts with blade 4 of RACK1 (91), sequence containing LARP4 CR1 and CR2, and ribosomal proteins, uS3, S16 and S17, that contact RACK1. The prediction resembled the RACK1-associated proteins as observed in the 80S ribosome with no apparent hindrance imposed by CR2 and CR1 (**Supp Fig S10B**).

### Advances in understanding LARP4 involvement in mRNA metabolism

The data in figure 3D and figure 4I suggest that the CR2-mediated RACK1-interaction is as protective as the PBM-mediated PABP-interaction in LARP4 stabilizing β-glo-ARE mRNA. The data are consistent with a model in which LARP4 promotes translation by stabilizing the ARE containing mRNA via 3’PAT protection while engaged by polysomes. Pruning was described to model deadenylation dynamics of subsets of long-lived, well-translated mRNAs in the context of their short PATs (43); thus, would be CNOT substrates while paradoxically remaining more refractory to decay than expected (7). For the abundant rp-mRNAs these appear as phased ∼30-nt mRNP-PABP-protected peaks in PATseq profiles, while long noncoding PAT-containing RNAs do not exhibit phasing (43). Although rp-mRNAs lack AREs or other elements that promote fast decay by CNOT, how they are maintained in a steady state of stabilization with short PATs is unclear though they may recruit pruning factors and PAT protection factors (2,7). Pruning proposes that stable mRNAs with short PATs, including closed loop mRNAs, exhibit high TE in a context of limited local PABP concentration and protected from deadenylation (43). Analysis of mRNA-PATseq-phasing in LARP4 knockout and control cells indicated protection of PABP-containing short-PAT rp-mRNPs from deadenylation and stabilization (15,22). LARP1-PABP protects short-PAT mRNPs from deadenylation, by the CNOT deadenylase globally, with greater effect on the rp-mRNAs (19). Thus, LARP1 and LARP4, and perhaps LARP1B and LARP4B would appear to be PAT protection factors for mRNA subsets including in a pruning model, consistent with limited local PABP concentration (18) (2,8,43). Yet, the extent to which PAT protection per se contributes to TE is unclear, and recruitment and/or retention of ribosomes or other activities may independently contribute (2,18). The results here indicate that the CR2 of LARP4 enables contact with RACK1 as an interaction bridge to ribosomes. They suggest that this promotes enhanced translational efficiency of mRNA in such context providing a basis for future studies.

## Abbreviations

Ab: antibody
ActD: actinomycin-D
b-glo: beta-globin
ARE: AU-rich element
EV: empty vector
GFP: green fluorescent protein
HEK293: human embryo kidney
PABP: poly(A)-binding protein
PAT: poly(A) tail
SM-PATseq: single molecule PAT sequencing
t½: half-life
TE: translational efficiency
WT: wild-type

## DATA AVAILABILITY

The data underlying this article are available in the article and in supplementary material.

## AUTHOR CONTRIBUTIONS

RM, SM, MRC, AR, ICG, JCC and LFP conceived experiments and analyzed data. RM, SM, MRC, AR and JCC wrote and/or edited the paper. RM conceived and oversaw the Y2H analysis. ICG performed ITC and physical association analyses with supervision from MRC. SM performed sequence analysis and designed mutagenesis for initial LARP4 *in vivo* IP analysis. AR performed experiments on LARP4 and mRNA expression, stability and polysome analysis. JCC performed *in vivo* experiments on LARP4B with supervision from MRC. LFP purified human LARP4 and RACK1 proteins for and performed pulldown assays. RM performed sequence and LOGO analyses as well as AFM predictions and analyses of RACK1 complex assembly of LARP4 and RACK1, RACK1-RPS3-RPS16-RPS17-LARP4, and RACK1-LARP4B, and graphics used for figure production.

## ACKNOWLEDGEMENTS

MRC and JCC thank Agi Grigoriadis for reagents, access to infrastructure and support. Authors thank the Centre for Biomolecular Spectroscopy (for ITC equipment) funded by the Wellcome Trust. RM thanks Eugene Valkov for sharing information on an AFM prediction of RACK1 complex with LARP4, suggestions for AFM prediction, computational analyses, and discussion, designing guided mutations, methods and reagents for pulldown experiments, supervision of LFP, and comments on an earlier version of the manuscript.

## SUPPLEMENTARY MATERIAL

### SUPPLEMENTARY FIGURE LEGENDS

**Supp Figure S1:**
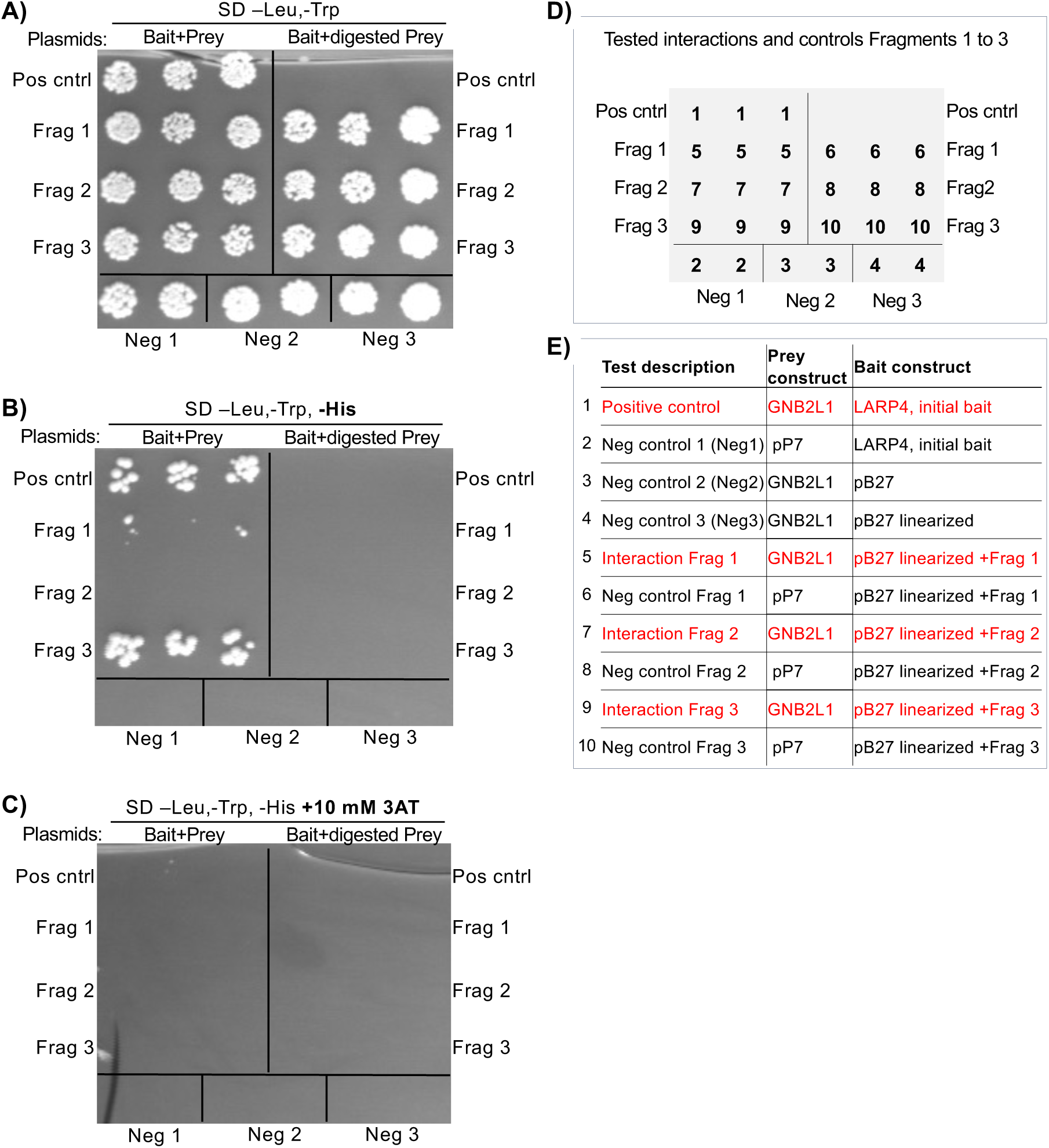
**A-C)** Yeast two hybrid domain mapping growth results according to plating layout design in panel **D)** with component constituents in panel **E)**. **A-C** are as in Figure 1, spotted cells grown from three isolated colonies on agar plates containing the selective media listed above. SD –Leu,-Trp selects for presence of the bait and prey plasmids respectively; SD –Leu,-Trp,-His additionally tests for interaction via production of the reporter gene product His3; and SD –Leu,-Trp,-His, **+**10 mM 3AT blocks function of the reporter product.

**Supp Figure S2:**
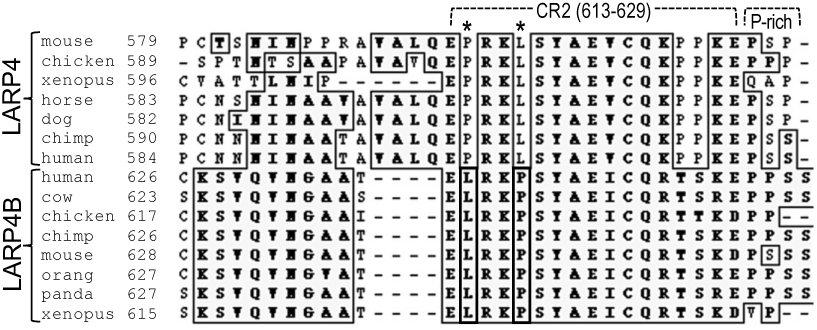
A) Multiple sequence alignment (MSA) of LARP4 and LARP4B. numbered above corresponding to conserved region 2 (CR2) of human LARP4, followed by bracket indicating a proline-rich region; asterisks above the sequence denote reciprocal/switched Leu and Pro in LARP4 and 4B.

**Supp Figure S3:**
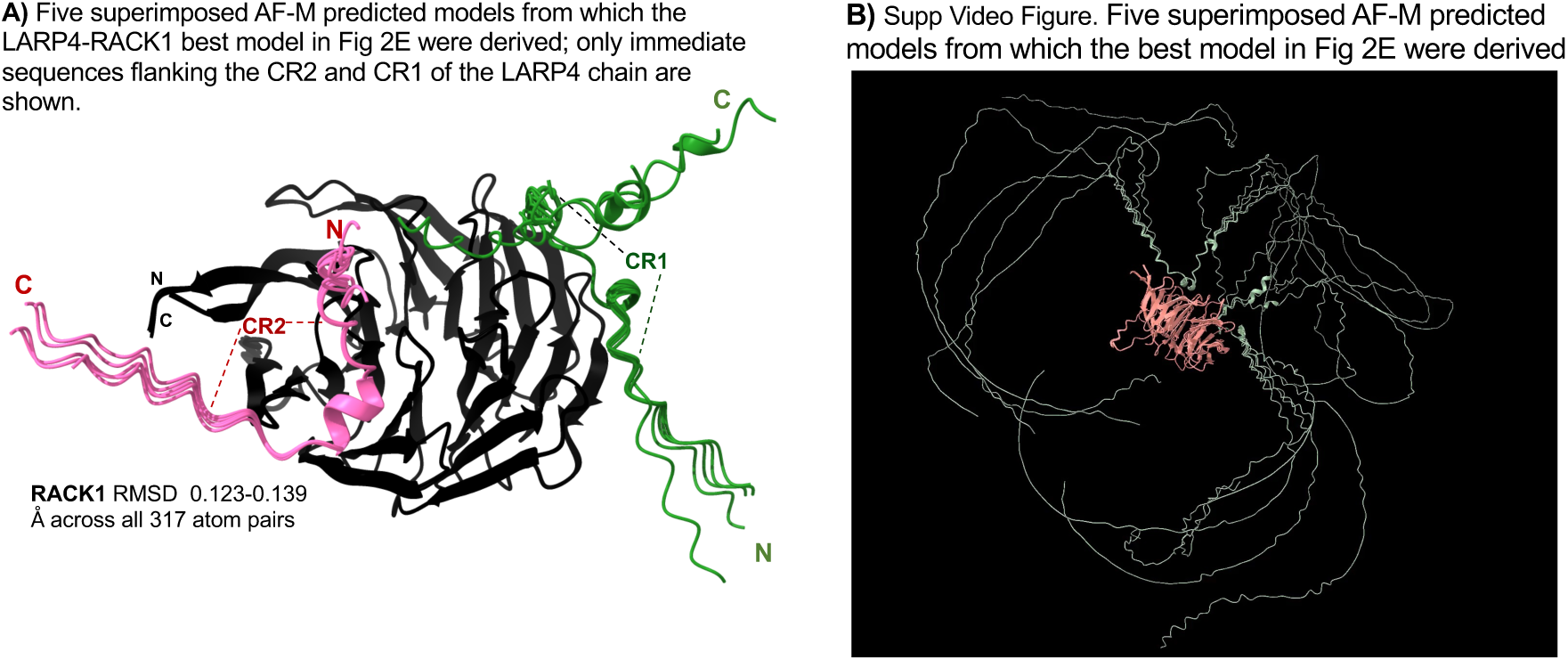
Five superimposed LARP4-RACK1 models predicted by AlphaFold-Multimer (AFM) corresponding to Fig 2E. The protein sequences input were LARP4(401–724) and RACK1 full length. **A)** LARP4 regions were rendered invisible except for those extending locally from CR1 and CR2 sequence motifs, the boundaries of which are indicated by dashed lines. RACK1 is black. ChimeraX Matchmaker analysis of the five AFM predictions revealed good agreement for RACK1 (RMSD across the 317 residue atom pairs whereas LARP4(401–724) agreement was limited to the CR1 and CR2 sequence regions. **B)** Video: see separate MP4 file. The five superimposed models predicted by AFM; as in A but with the views of LARP4 amino acids 401-724 visible.

**Supp Fig S4:**
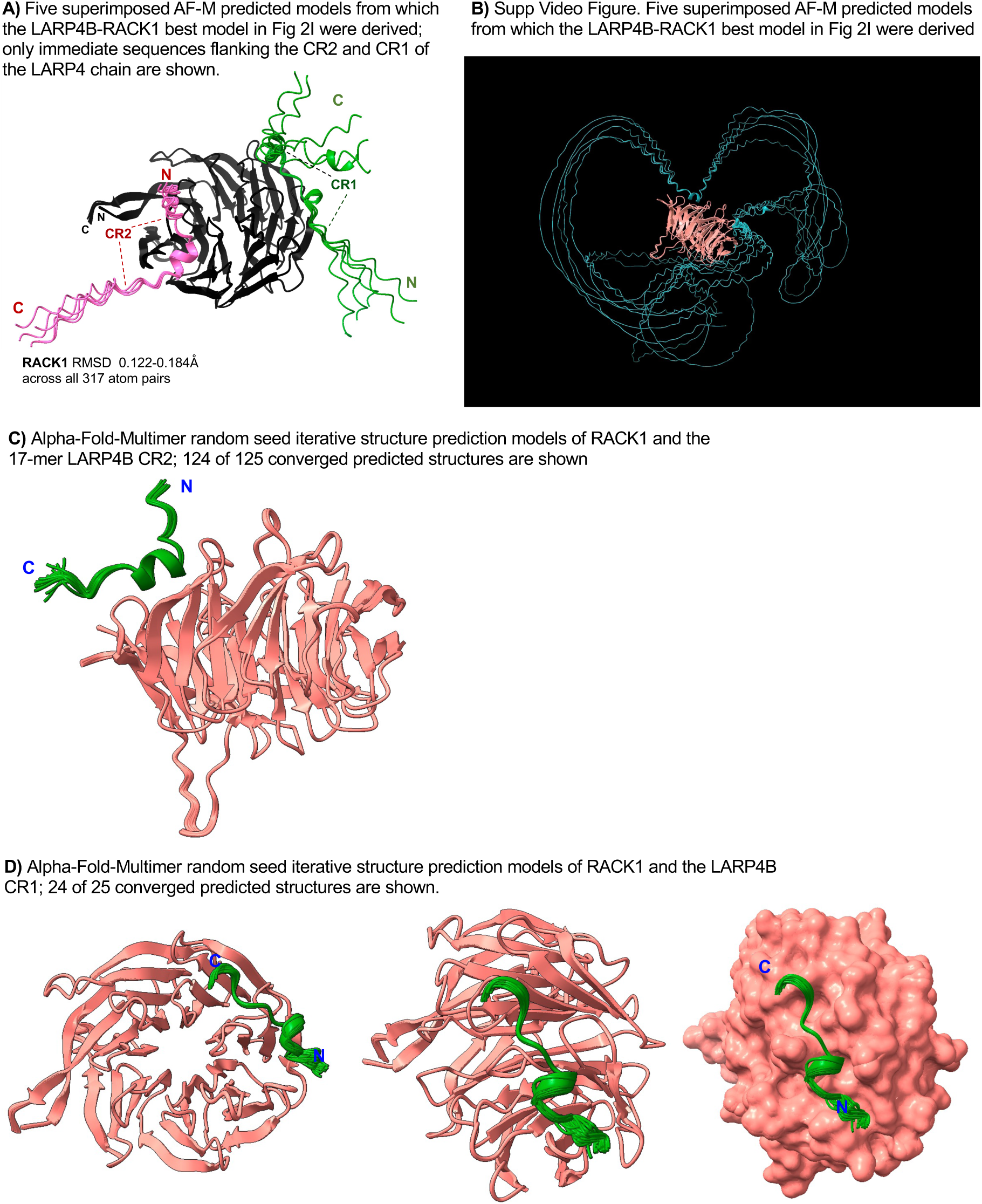
A) Five superimposed LARP4B-RACK1 models predicted by AFM corresponding to Fig 2J. The protein sequences input were LARP4B(433–738) and RACK1 full length. LARP4B regions were rendered invisible except for those extending locally from CR1 (green) and CR2 (pink) sequence motifs, RACK1 is black. ChimeraX Matchmaker analysis of the five predictions revealed good agreement for RACK1 (RMSD across the 317 residue atom pairs) whereas LARP4B(433–738) agreement was limited to the CR1 and CR2 sequence regions. **B)** Video: see separate MP4 file. The five superimposed AFM-predicted models as in A but with LARP4B amino acids 433-738 visible. **C-D)** Results of AFM random seed iterative model predictions. **C)** Protein sequence inputs were: RACK1 and only the 17-mer CR2 sequence, LARP4B(644–660). The figure shows converged predicted models (124 out of 125 models). **D)** Protein sequence inputs were: RACK1 and the 13-mer CR1 within a longer sequence, LARP4B(513–555); Left: RACK1 in ribbon view in orientation as in Fig 2I. Middle and right: rotated views.

**Supp Figure S5:**
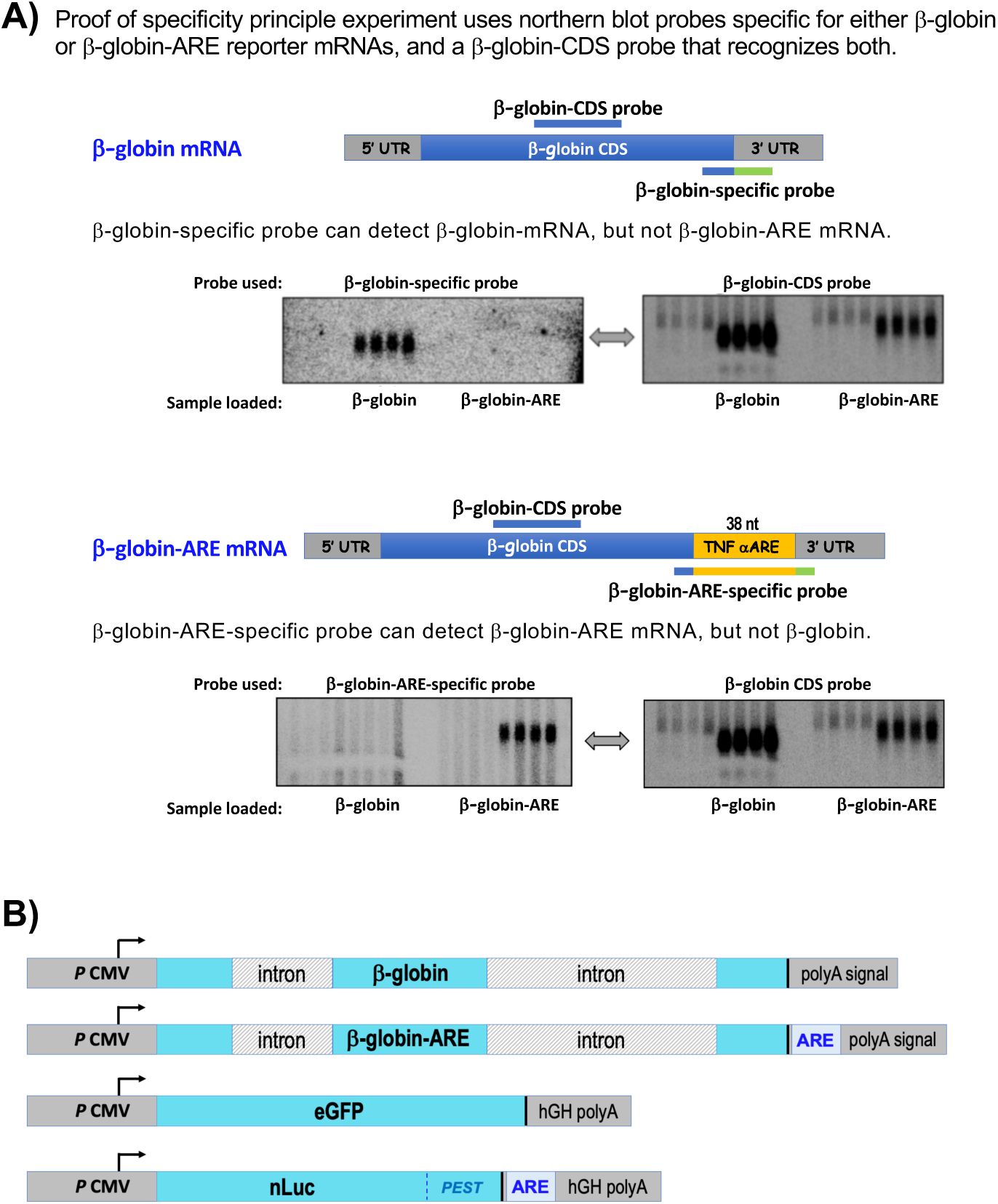
A) Northern blot oligo probes specific for β-Globin or β-Globin ARE reporter mRNAs corresponding to figures 3C and 4I. Schematic representation of the mature β-globin mRNA and β-globin-ARE mRNA reporters. The gene-specific probes used for the blots in the main figures are indicated by the 2 or 3 color bars below the cartoons. The β-globin-CDS probe indicated above the cartoons was used for the blots in this supplementary figure, for demonstrative purposes, as annotated. A single blot loaded with four lanes of β-glo mRNA on the left side and four lanes of β-globin-ARE mRNA on the right side, revealed all lanes by the β-globin-CDS probe. After stripping of the probe and subsequent reprobing, each gene-specific probe detected its target mRNA but not the other. **B)** Schematic representation of the plasmid reporter genes used in transient transfection experiments. Blue regions represent coding sequences; the stop codon is indicated by a thick vertical line. The β-globin and β-globin-ARE reporters differ only by the presence/absence of the 38 nt ARE in the 3’-UTR.

**Supp Figure S6:**
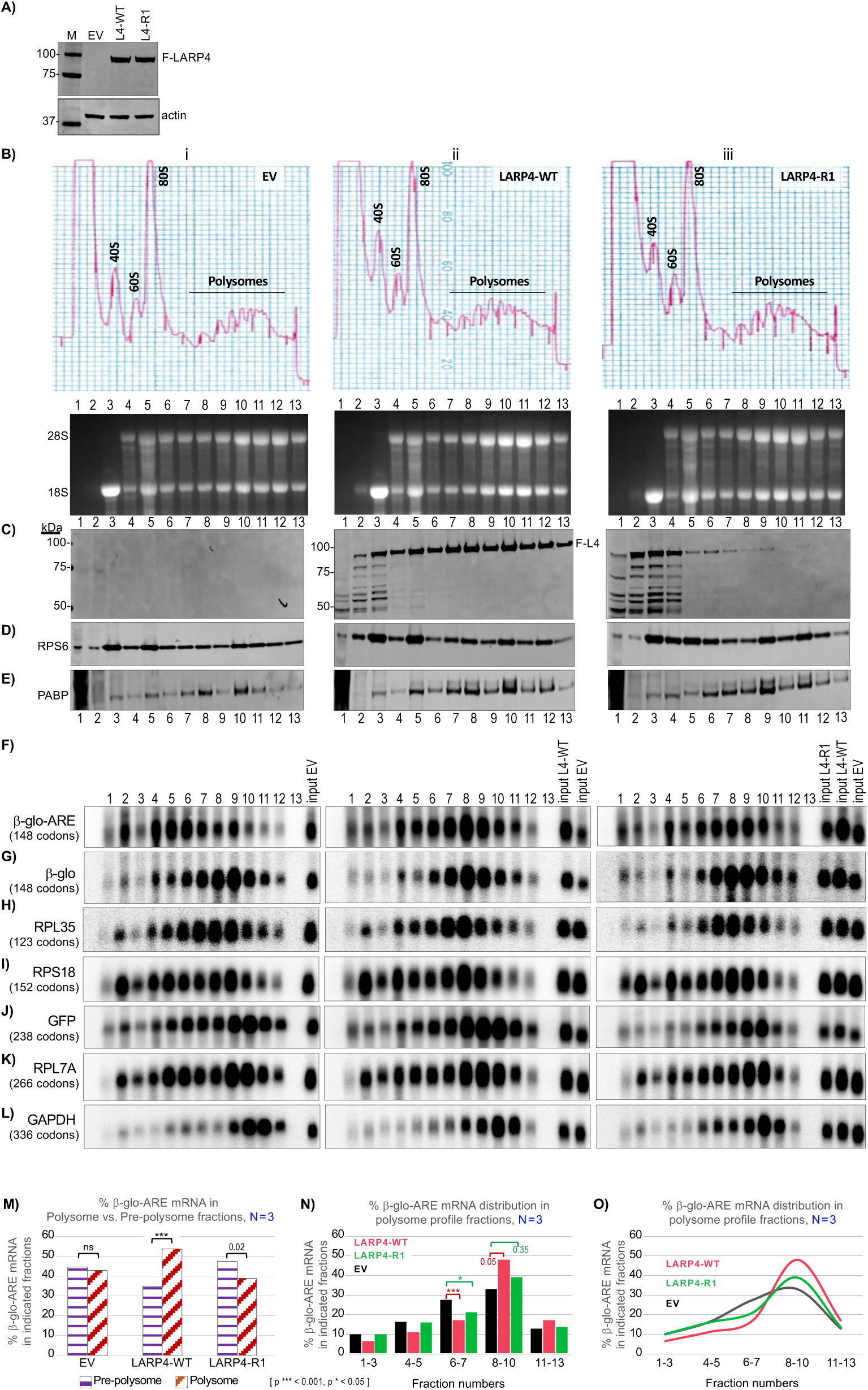
Replicate polysome profile data supporting figure 4, and quantitative analysis of β-globin-ARE mRNA distribution in polysome and pre-polysome fractions from three biological replicates. The layout is like figure 4. **A)** Immunoblot of input lysates from samples with transfected constructs indicated above the lanes, EV (empty vector), L4-WT and L4-R1 probed with anti-Flag Ab. **B)** Polysome profiles of the lysates run in parallel, representing EV, L4-WT and L4-R1 under which are an Ethidium Bromide-stained agarose gel, an immunoblot processed as indicated to the left (**C-E**), and a northern blot probed for various mRNAs (**F-L**), stacked in columns i-iii. **M-O)** Triplicate data set based on quantification of the distribution of β-glo-ARE mRNA among polysome profile fractions of three independent sucrose gradient sedimentations as in Fig 4, Supp Fig S5 and another. **M:** Graph of % β-glo-ARE mRNA in polysome vs. pre-polysome fractions; collective % in fractions 9-13 and in fractions 1-7 as indicated by bar patterns for EV, LARP4-WT and LARP4-R1. P value *** *p* <0.001 determined by two-tailed student’s T test; ns = non-significant. **N:** Triplicate β-glo-ARE mRNA polysome profile distribution in which amounts in each fraction set was plotted as bars. **O:** Same as in N, plotted as lines.

**Supp Figure S7:**
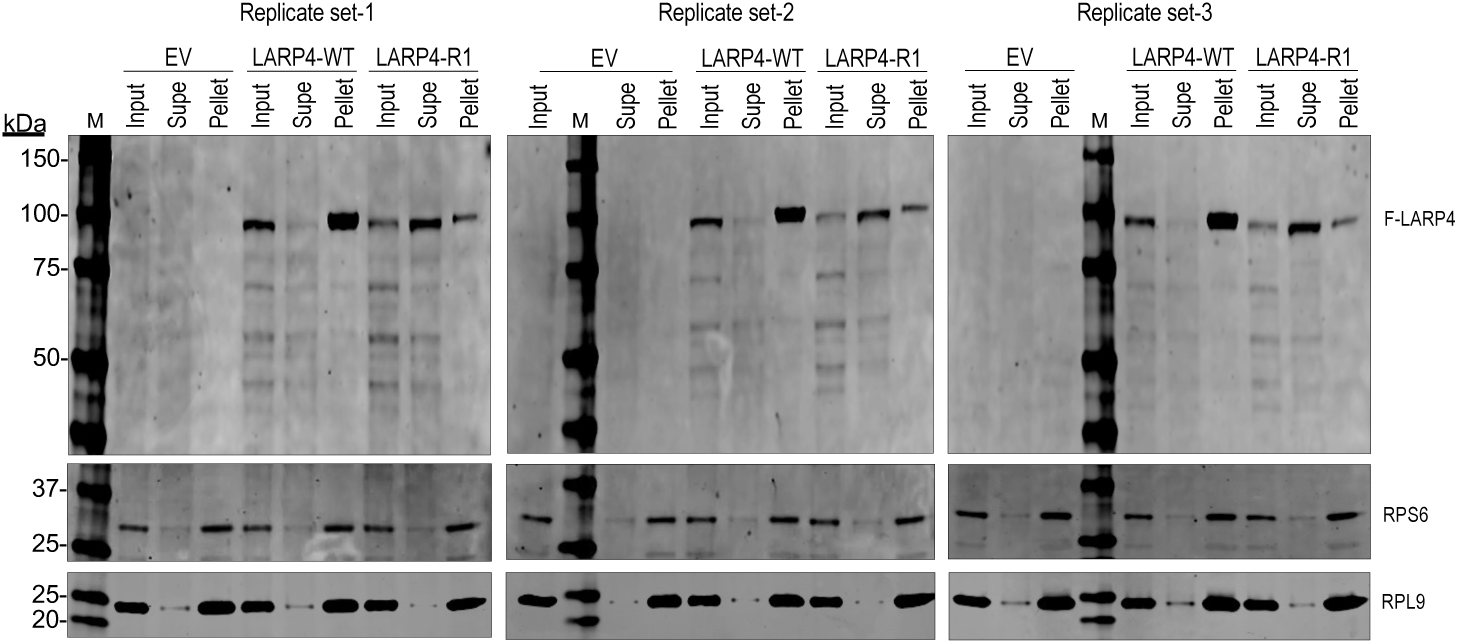
Analysis of LARP4 cosedimentation with ribosomes by S100 sucrose cushion; replicate data related to figure 4F. Here are the three replicate sucrose cushion S100 sedimentation analysis, each on an immunoblot processed for detection of Flag-L4, RPS6 and RPL9. The inputs (in), supernatants (S), and pellets (P) for the lysates from EV, LARP4-WT and LARP4-R1 samples are as indicated.

**Supp Figure S8:**
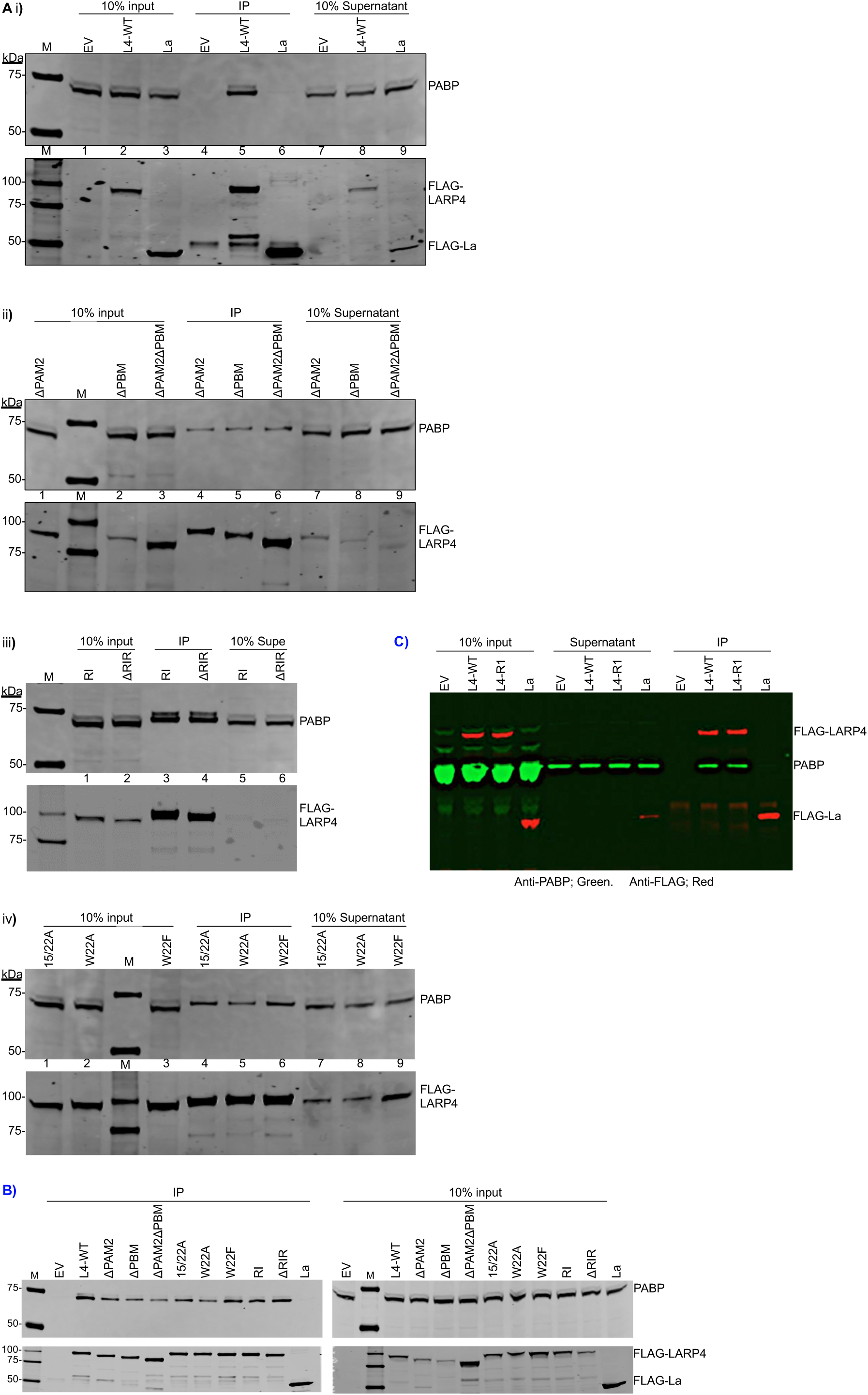
Co-IP of PABP with Flag-LARP4; replicate data sets related to figure 4H. **A)** Panels i, ii, iii and iv are immunoblot results of a single IP experiment in which all samples were processed side-by-side but too numerous to fit on one gel. The LARP4 mutated proteins indicated above the lanes were expressed as FLAG-tagged by transfection. Extracts were prepared and IPed using anti-Flag Ab. The blots in each panel show input extracts, IP products and supernatants, as indicated, processed for detection using anti-PABP (upper) and anti-Flag (lower). **B)** Independent experiment in which IP samples were on one blot and inputs on another; Flag-La was a negative control. **C)** An independent experiment in which only L4-WT, L4-R1 and La protein were compared for co-IP of PABP. The anti-Flag and anti-PABP are distinguished by color; Flag-La was a negative control.

**Supp Figure S9:**
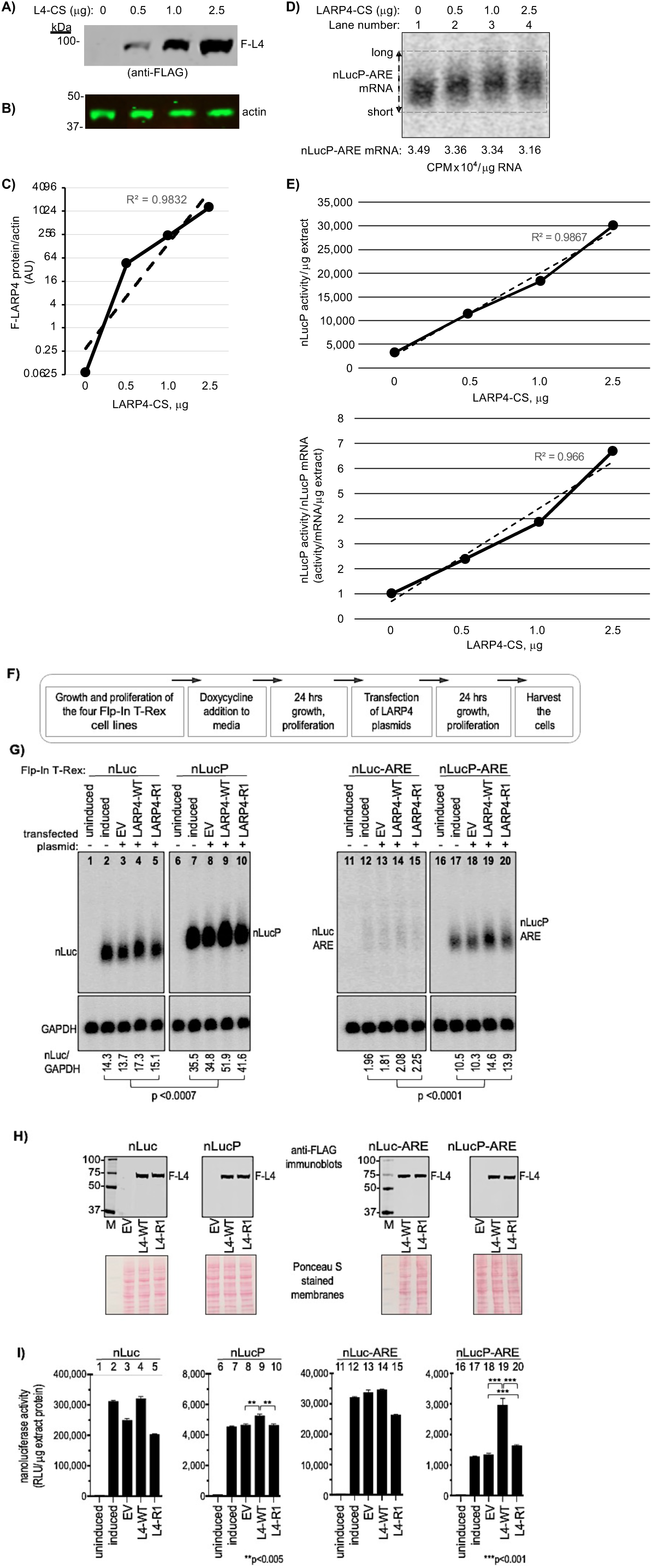
LARP4 activity increases translation efficiency of nanoLuciferase (nLuc) reporter mRNA. **A-E)** As in figure 5, 48 h after cotransfection of the nLucP-ARE reporter plasmid and the LARP4-CS (codon swap) expression plasmid, cells were harvested and half processed for protein and half for RNA. To each of the transfection reaction mixtures containing equal amounts of the nLucP-ARE reporter plasmid, was added the different amounts of LARP4-CS plasmid in μg indicated above the lanes (0 = empty plasmid). **A-C:** Analysis of LARP4-CS expression by immunoblot developed using anti-Flag (A), anti-actin (B), and the results were quantified and graphed (C). **D:** Northern blot analysis of nLucP-ARE mRNA expression in the same samples as in A-B as indicated above the lanes. The nLucP-ARE RNA reactive with the probe was quantified and indicated below the lanes. The double ended arrow at the left annotated with long and short refers to poly(A) length differences characteristic of LARP4 activity. A dashed line rectangle helps discern the lower and upper boundaries of that reflect poly(A) lengths that contribute to the mobility shift**^1-3^**. **E:** The upper graph plots nLucP luciferase activity in relative light units (RLU)/μg extract (Y-axis) versus LARP4-CS protein levels (X-axis). The R**^2^** value derived from a linear fit trendline is shown as a dashed line. The lower graph plots TE, (nLucP luciferase activity/μg extract)/(nLucP-ARE mRNA/μg extract) on the Y-axis versus LARP4-WT levels (X-axis). The R**^2^**value derived from a linear fit trendline is shown as a dashed line. **F-I)** Analysis of four nLuc reporters integrated in Flp-In HEK293 cells, and responses to LARP4-WT and LARP4-R1. **F:** Diagram showing experimental schema as in Figure 5. **G:** Northern blot analysis. The four panels contain RNAs from cells treated as indicated above the lanes, from the Flp-In T-Rex line containing the single copy reporter: nLuc, nLucP, nLuc-ARE and nLucP-ARE. All samples were processed side-by-side; the two left and two right panels were run on different gels and transferred to different blots which were hybridized with the same antisense nLuc probe, washed together, and exposed on the same phosphorImager screen. Quantification of the nLuc signal/GAPDH is shown below the lanes. **H:** Immunoblot analysis of the Flag-LARP4 levels as indicated. **I:** Nanoluciferase activities in the protein extracts corresponding to the samples in G; note different scales on Y-axes. **J:** Translation efficiency (TE). The nanoluciferase activity data from I was normalized by the corresponding nLuc-reporter mRNA and plotted.

**Supp Figure S10:**
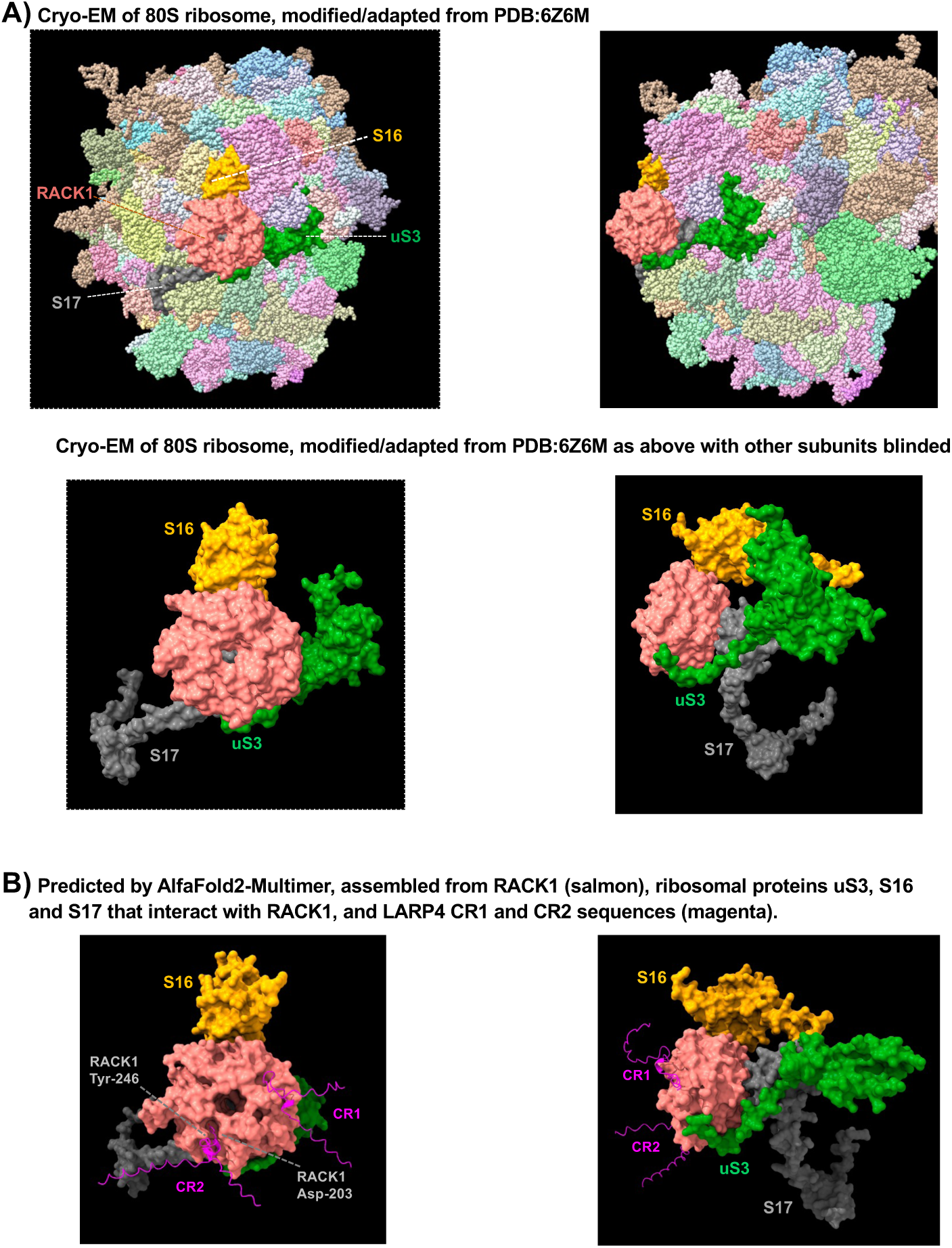
**A)** Views of RACK1 as it resides on an existing structure of the human 80S ribosome (PBD: 6Z6M**^4^** ^see^ **^5^**. The upper structure was oriented such that RACK1 is central and facing the viewer; it and the three native ribosomal proteins in close contact, RPS3, RPS17, and RPS6, were rendered in surface view as indicated. For the bottom panel, all elements in PBD 6Z6M other than the four shown above in surface view were made invisible, and features of RACK1 were annotated (see Fig 2E). **B)** AlphaFold-Multimer was used in standard mode to predict a model assembled from RACK1, and sequences representing LARP4, RPS3, RPS17, and RPS6 (Methods), sequence coverage for each component was >1000. LARP4 residues were rendered invisible except for those extending locally from the CR1 and CR2 sequences. The best model is shown. AFM produced a predicted assembly similar in general orientation to that in the lower panel of A. In B, the CR2 and CR1 motifs are engaged in much the same way as for RACK1 alone.

**Supp Figure S11:**
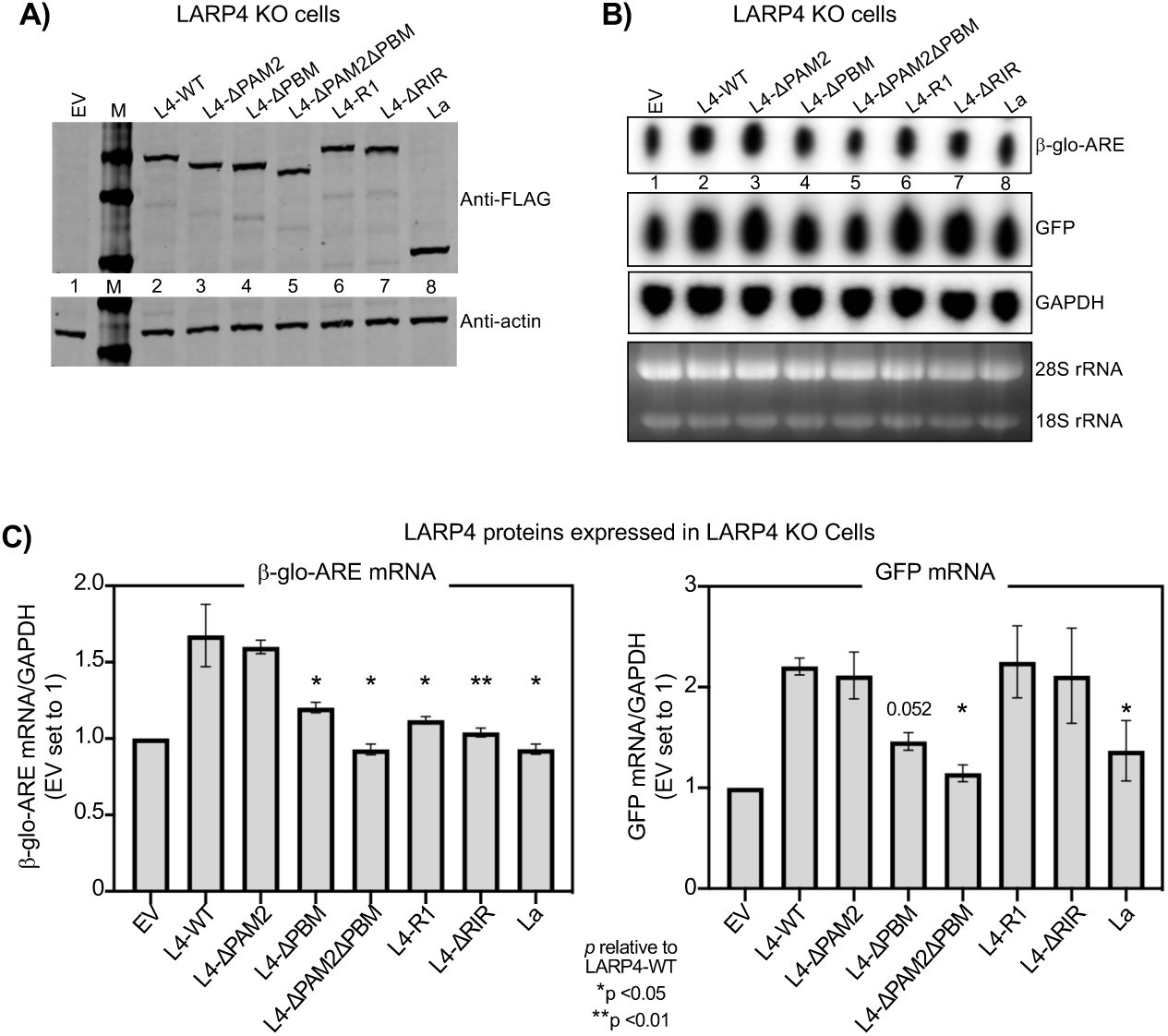
β-glo-ARE mRNA stabilization in LARP4-KO cells is rescued differentially by LARP4-WT and LARP4-RACK1 mutants. **A)** Immunoblot of extract protein from HEK293 LARP4-KO cells transfected with the LARP4 (L4) constructs above the lanes; EV: empty vector (see text). **B)** Northern blot shows simultaneous responses of β-glo-ARE and GFP mRNAs in the same pool of transfected cells carrying a different LARP4 mutant. **C)** Quantitation of northern blot signals of biological duplicate experiments. Transcript levels were normalized by GAPDH mRNA.

